# Light-dependent translation change of Arabidopsis *psbA* correlates with RNA structure alterations at the translation initiation region

**DOI:** 10.1101/2020.06.11.145870

**Authors:** Piotr Gawroński, Christel Enroth, Peter Kindgren, Sebastian Marquardt, Stanisław Karpiński, Dario Leister, Poul Erik Jensen, Jeppe Vinther, Lars B. Scharff

**Author notes:** These authors contributed equally to this work (Piotr Gawroński, Christel Enroth). Correspondence to: Lars Scharff.

## Abstract

mRNA secondary structure influences translation. Proteins that modulate the mRNA secondary structure around the translation initiation region may regulate translation in plastids. To test this hypothesis, we exposed *Arabidopsis thaliana* to high light, which induces translation of *psbA* mRNA encoding the D1 subunit of photosystem II. We assayed translation by ribosome profiling and applied two complementary methods to analyze *in vivo* RNA secondary structure: DMS-MaPseq and SHAPE-seq. We detected increased accessibility of the translation initiation region of *psbA* after high light treatment, likely contributing to the observed increase in translation by facilitating translation initiation. Furthermore, we identified the footprint of a putative regulatory protein in the 5’ UTR of *psbA* at a position where occlusion of the nucleotide sequence would cause the structure of the translation initiation region to open up, thereby facilitating ribosome access. Moreover, we show that other plastid genes with weak Shine-Dalgarno sequences (SD) are likely to exhibit *psbA*-like regulation, while those with strong SDs do not. This supports the idea that changes in mRNA secondary structure might represent a general mechanism for translational regulation of *psbA* and other plastid genes.

**SIGNIFICANCE:** RNA structure changes in the translation initiation region, most likely as a result of protein binding, affect the translation of *psbA* and possibly other plastid genes with weak Shine-Dalgarno sequences.

## INTRODUCTION

The secondary structure of mRNA is important for many translation related processes in bacteria and bacteria-derived eukaryotic organelles. This includes the efficiency of translation initiation (de Smit and van Duin, 1990; Kudla et al., 2009; Goodman et al., 2013; Mustoe et al., 2018; Bhattacharyya et al., 2018), the recognition of start codons (Scharff et al., 2011; Nakagawa et al., 2017; Scharff et al., 2017), and ribosome pausing (Wen et al., 2008; Tuller et al., 2011; Gawroński et al., 2018). In addition, changes in mRNA secondary structure can regulate translation initiation. Some of the mechanisms involved, such as riboswitches (Breaker, 2018) and RNA thermometers (Neupert et al., 2008; Krajewski and Narberhaus, 2014), are independent of proteins, whereas others depend on the binding of small RNAs or proteins to either activate or repress translation by modifying mRNA secondary structure (Laursen and Sørensen, 2005; Duval et al., 2015).

In plastids – plant organelles derived from cyanobacteria – changes in mRNA secondary structure have also been proposed to regulate translation (Stampacchia et al., 1997; Klinkert et al., 2006; Prikryl et al., 2011; Hammani et al., 2012). This is not surprising as in plastids, bacterial-type 70S ribosomes synthesize proteins and the process shows many similarities to translation in bacteria. Indeed, translational regulation is a major determinant of gene expression in plastids (Barkan, 2011; Sugiura, 2014; Sun and Zerges, 2015; Zoschke and Bock, 2018). The intrinsic mRNA features that determine the efficiency of start codon recognition in plastids of higher plants, and hence the efficiency of translation initiation, are well characterized: a) Shine-Dalgarno sequences hybridize to the anti-Shine-Dalgarno sequence at the tail of the 16S rRNA and thereby position the start codon so that it can bind to the initiator tRNA; b) local minima of mRNA secondary structure around the start codon make it accessible for the ribosome, whereas other AUGs are masked by folded RNA (Hirose and Sugiura, 2004; Scharff et al., 2011; Zhang et al., 2012; Scharff et al., 2017; Gawroński et al., 2020).

Compared to the intrinsic mRNA features determining the efficiency of translation initiation, we understand much less about the molecular mechanisms regulating translation in plastids. One hypothesis is based on *in vitro* findings that some RNA-binding proteins can alter the structure of the translation initiation region of their target mRNAs in a way which activates translation initiation (Stampacchia et al., 1997; Klinkert et al., 2006; Prikryl et al., 2011; Hammani et al., 2012). In the absence of such a protein, the Shine-Dalgarno sequence and/or the start codon are occluded by mRNA secondary structure; therefore, translation efficiency is low. The binding of the regulatory protein shifts the structural equilibrium to an RNA conformation that makes these *cis*-elements, which are essential for translation initiation, accessible to the ribosome and thereby activates or upregulates translation.

Here, we tested this hypothesis by exposing *Arabidopsis thaliana* plants to high light. In higher plants, this condition is known to induce the translation of the plastid-encoded *psbA* mRNA (encoding the D1 subunit of photosystem II) on the level of translation initiation (Chotewutmontri and Barkan, 2018; Schuster et al., 2020). This increase of *psbA* translation counteracts the increase in D1 turnover due to photodamage (Mulo et al., 2012; Li et al., 2018). Changes in the *psbA* mRNA *in-vivo* secondary structure and translation efficiency were analyzed, and our findings support that *psbA* is regulated by an RNA-binding protein that increases the accessibility of the Shine-Dalgarno sequence under high light conditions. Moreover, our analysis of the relationship between mRNA secondary structure and translation suggests that this mechanism is generally used to regulate translation of plastid-encoded genes that, like *psbA*, possess a weak Shine-Dalgarno sequence.

## RESULTS

We tested whether changes in mRNA secondary structure could influence translation in chloroplasts using a well-known example for translation regulation as a starting point: the high light induced upregulation of *psbA* translation, the mRNA coding for the D1 subunit of photosystem II (Mulo et al., 2012; Schuster et al., 2020). First, we validated that in young *Arabidopsis thaliana* plants (17-18 days old) *psbA* translation was induced by exposure to high light for one hour. For this, we extracted polysomes, size-fractionated them in sucrose gradients, and analyzed the distribution of *psbA* mRNA by RNA gel blot analysis. As expected, we observed a prominent shift of *psbA* mRNA into denser fractions (relative to low light controls), which indicates increased loading of ribosomes and higher translation initiation rates in high light (Supplemental Figure S1).

### mRNA secondary structure changes in the *psbA* translation initiation region

Next, we focused our analysis on the translation initiation region of the *psbA* mRNA and analyzed its *in-vivo* secondary structure using dimethyl sulfate (DMS) probing. DMS was described to methylate only N1 of adenosines and N3 of cytidines of single-stranded and accessible RNA (Mitchell et al., 2019). However, recently, it was demonstrated that under alkaline conditions DMS can probe also guanosines and uridines (Mustoe et al., 2019). As the chloroplast stroma is slightly alkaline, all four nucleotides of chloroplast RNA can be probed, although the probing of adenosines and cytidines is more reliable (Gawroński et al., 2020). High DMS probing at a nucleotide indicates a single-stranded confirmation. Low probing can be caused by double stranded regions, protein binding or compact RNA secondary structure preventing DMS access (Mitchell et al., 2019). DMS efficiently enters cells (Wells et al., 2000), including those of Arabidopsis plants (Ding et al., 2014), and is therefore suited for *in-vivo* structural probing. DMS-reactivity of probed nucleotides can be quantified by mutational profiling (MaP) using a thermostable group II reverse transcriptase (TGIRT), which during reverse transcription incorporates mutations in the cDNA at the reacted positions (Zubradt et al., 2017). The young plants were exposed to either low light or high light for one hour and then, in the same light regime, incubated in a DMS solution for six minutes (see Methods section for details). The probing did not cause browning of the leaves as previously observed (Wang et al., 2019), and the quality of the extracted RNA was not affected by the treatment (Supplemental Figure S2). Using gene-specific primers, we analyzed the translation initiation region of *psbA* and, as a control, helix 33 of the plastid 16S rRNA (see Supplemental Table 1 for the coverage). In parallel, we probed purified RNA that had been refolded *in vitro* (Figure 1, Supplemental Figure S3, S4, S5). In addition, we analyzed young plants treated with water instead of DMS under low light and high light conditions and found a very low background level of mutations using our protocol (Supplemental Figure S3A, S4, S5B). As expected for DMS, adenosines and cytidines are statistically significantly more probed than guanosines and uridines (Supplemental Figure S3A). Furthermore, the observed DMS probing for helix 33 of the 16S rRNA corresponds nicely with the rRNA structure previously described for plastid ribosomes (Ahmed et al., 2017) (Supplemental Figure S5) and is similar for low and high light conditions (Supplemental Figure S3C, S5B). The structure signals for guanosines and uridines *in vivo* were, as expected (Mustoe et al., 2019; Gawroński et al., 2020), weaker than those for adenosines and cytidines, but still informative compared to the *in vitro*, protein-free control (Supplemental Figure S5A). In addition, the reproducibility of the probing of adenosines and cytidines was better than for guanosines and uridines (Supplemental Figure S3B).

**Figure 1.**
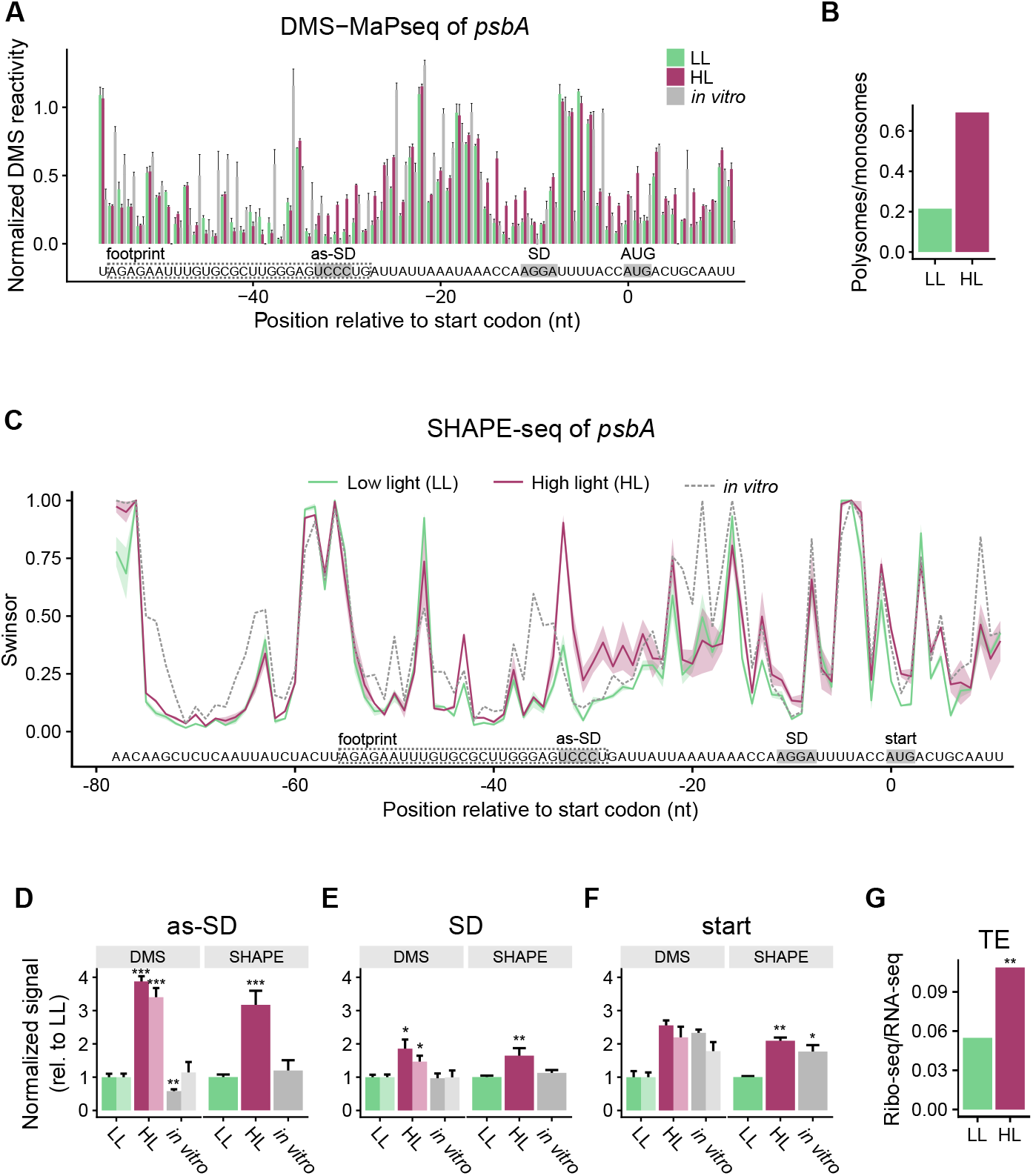
mRNA secondary structure changes in the translation initiation region of *psbA*: Increased accessibility of the Shine-Dalgarno sequence and the start codon correlates with increased translation efficiency. **A**, DMS-based mapping of *psbA* mRNA secondary structure of young, 17-18-days-old plants as determined by MaPseq. Normalized DMS reactivities are given to account for the differences between adenosines/cytidines vs. guanosines/uridines reactivities (Supplemental Figures S3A and S4A). Furthermore, the values for control without added DMS (Supplemental Figure S4A) were subtracted and outliers were removed by winsorization (only the 90th percentile is retained). The information obtained at adenosines/cytidines is more reliable than at guanosines/uridines (Supplemental Figures S3B and S5A) (Gawroński et al., 2020). High normalized DMS reactivity indicates single-stranded nucleotides. The data for the low light (LL) control are shown in light green, the high light (HL) samples in dark red, and the mRNA that was allowed to fold *in vitro* in gray. The error bars indicate the mean standard error. The start codon (start), Shine-Dalgarno sequence (SD), and a sequence that can bind the SD (as-SD) are marked. The position of the footprint of a putative regulatory protein is given as a dashed box (Supplemental Figure S11A). A comparison of the DMS-probed RNA with a water-treated control is shown in Supplemental Figure S4A. **B**, Polysome analysis of *psbA* translation in 17-18-days-old plants (Supplemental Figure S1). The ratio of the *psbA* mRNA **C**, SHAPE analysis (NAI-N_3_ probing) of young leaves of 7-week-old plants. SHAPE signals indicate the extent to which each nucleotide is unpaired. Swinsor values are the termination counts, i.e. how often reverse transcription was stopped at each nucleotide by a bound NAI-N_3_ probe, normalized by winsorization (only the 90th percentile is retained, outliers are discarded). High swinsor values indicate unpaired nucleotides; low swinsor values base-paired nucleotides. The shaded areas around the lines indicate the mean standard error. **D**, Average of mRNA secondary structure at the sequence binding the Shine-Dalgarno sequence (as-SD), as revealed by DMS and SHAPE values normalized to the low light values. The columns in darker color represent the more reliable reactivities at adenosines/cytidines, the lighter the reactivities at all four nucleotides. Asterisks here (and in **E** and **F**) indicate statistically significant changes compared to LL (*P*-values calculated with the Wilcoxon rank sum test; * = *P* < 0.05, ** = *P* < 0.01, and *** = *P* < 0.001), error bars indicate mean standard error. **E**, Average structure at the Shine-Dalgarno sequence (SD). **F**, Average structure at the start codon (start). **G**, Change in translation efficiency (ratio footprints/transcript levels) of *psbA* mRNA in young leaves of 7-week-old plants (Figures 4, S12, and S13). Asterisks indicate statistically significant changes (calculated with RiboDiff; ** = *P* < 0.01).

The DMS-MaPseq results for the *psbA* translation initiation region were highly reproducible (Supplemental Figure S3B,C,D). Obvious differences were detected around the Shine-Dalgarno sequence and the start codon (Figure 1A,E,F). In high light, both elements had higher DMS probing than in the low light samples. This was true when the probing of all four nucleotides was considered as well as when only the more reliable data at adenosines was considered (Figure 1E,F). The increased DMS probing indicates that these RNA regions are more single-stranded and accessible under high light conditions, which is in agreement with the observed increase of *psbA* translation (Figure 1B, Supplemental Figure S1). Interestingly, an upstream sequence, complementary to the Shine-Dalgarno sequence, also displayed increased DMS probing under high light conditions, suggesting that this sequence might interact with the Shine-Dalgarno sequence under low but not under high light conditions, and thus could control translational activation (Figure 1A,D). This would be in agreement with the hypothesis that translation efficiency is low when the Shine-Dalgarno sequence and/or the start codon are occluded in a double-stranded region. The opening of the structure would make these elements more accessible, which should boost translation efficiency.

To further validate the observed structural changes in the *psbA* mRNA, we used a complementary method, selective 2’-hydroxyl acylation analyzed by primer extension (SHAPE) (Spitale et al., 2013, 2014), to probe the RNA structure. Furthermore, to test the robustness of the response, we applied SHAPE to analyze *psbA* secondary structure changes in response to high light acclimation in mature plants. Arabidopsis plants grown in short-day, low light conditions were acclimated to high light by exposing 7-week-old plants to four hours high light, 16 hours dark, and again four hours high light. To analyze translation, we used young leaves, which were found to be more capable of acclimating to the high light conditions than mature, fully expanded leaves (Supplemental Figure S6). In agreement with the results obtained for the long day-grown young plants (Supplemental Figure S1), *psbA* translation was increased after one hour in high light (Supplemental Figure S7). This increase was still present after one day in the above-described acclimation conditions (Supplemental Figure S7). For RNA secondary structure probing, we analyzed RNA after one day high light acclimation using the SHAPE reagent NAI-N_3_ (Spitale et al., 2013, 2014). Like other SHAPE reagents, NAI-N_3_ reacts with the 2′-hydroxyl groups in the RNA backbone when the RNA adopts specific conformations that are characteristic for flexible single-stranded RNA, but it does so much less efficiently if their flexibility is constrained by base pairing (Merino et al., 2005; McGinnis et al., 2012). Hence, NAI-N_3_ effectively probes for the presence of single-stranded nucleotides. The SHAPE reactivity profile can be read out by mapping termination sites of reverse transcription caused by the introduction of SHAPE adducts, and NAI-N_3_ can be used for intracellular probing experiments (Spitale et al., 2013). We added NAI-N_3_ to flash-frozen leaf samples and probed the RNA during the thawing of the high light and low light samples. In addition, we performed *in-vitro* probing on purified RNA, which had been refolded *in vitro*. We performed SHAPE selection on the samples, as previously described (Poulsen et al., 2015), and the counts obtained were normalized for coverage using Smooth Winsorization (Kielpinski et al., 2015) to give SHAPE reactivities between 0 and 1.

First, we investigated the correlation between the replicates in our samples using PCA analysis (Supplemental figure S8A). As expected, the quality of the probing signal was dependent on the sequence coverage; therefore, we limited our analysis to RNAs having on average more than 10 termination counts per nucleotide (Supplemental figure S8B). In the PCA plot, the *in-vitro* probing data is clearly separated from the *in-vivo* samples, and the two high light samples cluster together. Among the five low light samples, three samples clustered together, whereas the remaining two deviated both from the other three and from each other; therefore, we excluded these two samples from our further analysis. Next, we checked the structural signal in the dataset by comparing the SHAPE probing data for the Arabidopsis 18S rRNA with the known secondary structure of this RNA. For all samples, except those having low coverage of the 18S rRNA owing to prior rRNA depletion, we observed a signal for RNA structure (Supplemental Figure S9).

For the translation initiation region of *psbA*, we observed good reproducibility of SHAPE reactivities among replicates (Figure 1C). The Shine-Dalgarno sequence, start codon, and the sequence that can potentially bind the Shine-Dalgarno sequence (as-SD), showed higher SHAPE reactivity in high light samples than in low light controls (Figure 1C-F). The SHAPE reactivities correlate with the DMS-MaP signal observed in the translation initiation region, especially with the more reliable DMS probing of adenosines and cytidines in the region from the as-SD to the start codon (Figure S10). Thus, using two different chemical probes, we find that the *psbA* translation initiation region becomes more accessible under high light conditions (Figure 1A,C,E,F, Supplemental Figure S10). The effect correlates well with increased *psbA* translation (Figure 1B,G) and is observed after both short-term high light stress in young plants and long-term high light acclimation of young leaves of 7-week-old plants.

### Change of mRNA secondary structure of *psbA* translation initiation region likely caused by protein binding

One potential means of altering mRNA secondary structure is the binding of RNA-binding proteins. Analyzing the reads from MNase-digested RNA, we detected a footprint of a putative regulatory protein in the 5’ UTR of *psbA* (Supplemental Figure S11A). We confirmed the footprint by northern blot analysis of small RNAs isolated without prior RNase treatment using a probe specific for the footprint sequence (Supplemental Figure 11D) as it was done previously (Loizeau et al., 2014; Ruwe et al., 2016). A central part of this footprint has previously been described as a site where HCF173 binds alone or with other unknown proteins (McDermott et al., 2019). HCF173 is a protein that activates *psbA* translation (Schult et al., 2007; Link et al., 2012). The detected footprint is located upstream of the Shine-Dalgarno sequence. We assessed the possible influence of a bound protein on the *psbA* mRNA secondary structure by predicting the structure using DMS reactivities and the position of the bound protein as constrains. The prediction revealed that the footprint contains sequences that can bind to the Shine-Dalgarno sequence and the start codon. In low light, these *cis*-elements are part of a double-stranded structure (Figure 2A). In high light, the Shine-Dalgarno sequence and the start codon are in a largely single-stranded structure and therefore more accessible (Figure 2B). The abundance of the footprint increased in high light compared to low light (Supplemental Figure S11A,B,D). However, we also observed increased *psbA* mRNA levels under high light conditions (Supplemental Figure S11C), and this could potentially explain the increased accumulation of small RNAs stemming from this region.

**Figure 2.**
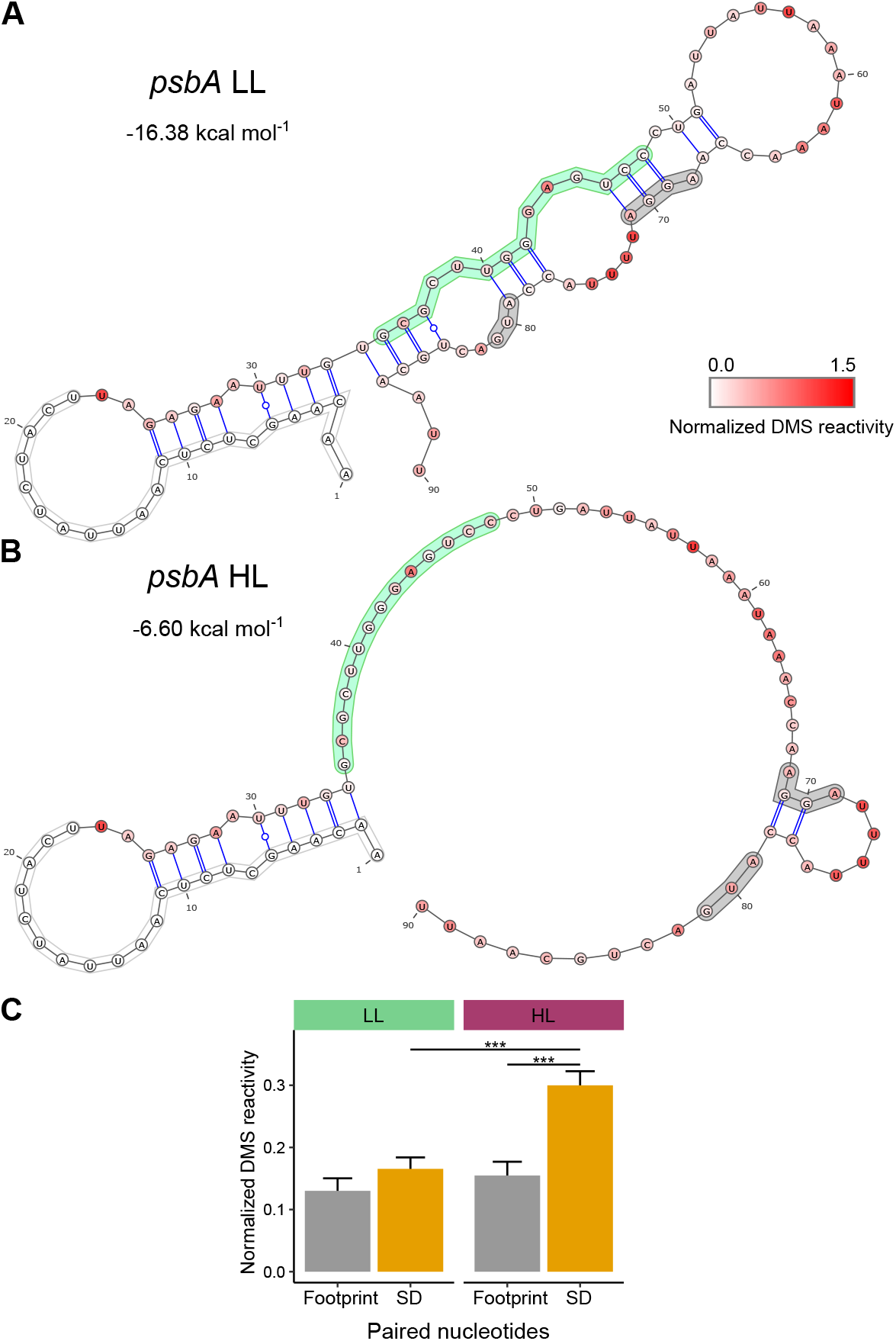
Footprint of a putative regulatory protein bound to the 5’ UTR of *psbA*: **A**, Predicted mRNA secondary structure of the *psbA* translation initiation region in low light (LL) using normalized DMS reactivities (Figure 1A) as constrains. The white box marks the position of the primer used to amplify the cDNA. For this region no DMS reactivities could be obtained. The green box marks the footprint of a RNA binding protein (McDermott et al., 2019) (Supplemental Figure S11A-D), the grey boxes indicate the Shine-Dalgarno sequence (AGGA) and the start codon (AUG). For each nucleotide, the normalized DMS reactivity is shown in a color code. The kcal mol^−1^ value for the strength of the RNA structure is given. **B**, Predicted mRNA secondary structure in high light (HL) using normalized DMS reactivities (Figure 1A) and the protein binding site (forced to be single-stranded) as constrains. For the structure predictions for *in vitro*-folded RNA see Supplemental Figure S11G. **C**, Normalized DMS reactivities of the nucleotides predicted to form base pairs in low light (**A**) between the region of the footprint (between nucleotides 35-48) and the region including the start codon and the Shine-Dalgarno sequence (SD) (between nucleotides 69-86). The average normalized DMS reactivities are shown separately for both regions. Nucleotides in these regions predicted not to be paired are excluded. DMS reactivities at the SD side significantly increase in high light indicating a shift to single-stranded RNA. This suggests that in low light a stem loop structure is formed (**A**), whereas in high light a protein is bound to the *psbA* translation initiation region making the SD and the start codon accessible (**B**). Asterisks indicate statistically significant changes compared to LL (*P*-values calculated with the Wilcoxon rank sum test; *** = *P* < 0.001), error bars indicate mean standard error. For the separately analyzed DMS reactivities at adenosines and cytidines as well as SHAPE reactivities see Supplemental Figure S11E,F.

As an alternative approach to distinguish between double-stranded RNA and a bound protein, we analyzed the DMS reactivity at the nucleotides of the footprint and around the Shine-Dalgarno sequence that are predicted to pair in low light (Figure 2A). DMS probing is sensitive to protein binding (Kwok et al., 2013; Talkish et al., 2014), therefore DMS reactivity is low both for a nucleotide bound to a protein and a nucleotide involved in base pairing. High DMS reactivity indicates a single-stranded, not protein-bound nucleotide. The half of the stem loop to which the protein binds had low DMS reactivities both in low light and high light (Figure 2C). In contrast, the DMS reactivities of the other half of the stem loop, at the sequence around the Shine-Dalgarno sequence and the start codon, increased in high light (Figure 2C). This suggests that these nucleotides pair in low light to nucleotides of the protein-binding site. In high light, a protein prevents the formation of the double-stranded structure and thereby increases the accessibility of the *cis*-elements required for translation initiation and *psbA* translation. The analysis of DMS reactivities only at adenosines and cytidines showed the same trend as the one for the DMS reactivities at all four nucleotides (Supplemental Figure S11E). Interestingly, whereas the average DMS reactivities of paired nucleotides at the footprint were similar in low light and high light (Figure 2C), the DMS reactivities of the nucleotides that can bind the Shine-Dalgarno sequence increased in high light (Figure 1A,D). This sequence is located at the 3’ end of the footprint (Figure 1A) but is still part of the footprint, which is protected from nuclease attack. A possible explanation is that the protein binding at the footprint interacts only with some of the nucleotides and therefore does not influence the DMS reactivities of the other nucleotides (compare also Figure S10). The SHAPE reactivities differ from the DMS reactivities (Supplemental Figure S11F), which can be caused by differences between SHAPE reagents and DMS in the sensitivity to protein binding. DMS reactivity is low in case of bound proteins (Kwok et al., 2013; Talkish et al., 2014), whereas bound proteins are not always detected as nucleotides with low SHAPE reactivity (Spitale et al., 2013, 2015; Kenyon et al., 2015) (compare also Figure S10).

### mRNA secondary structure of the translation initiation regions of *rbcL*

As an additional example, we examined the translation initiation region of *rbcL*, which encodes the large subunit of RuBisCO. In contrast to *Nicotiana tabacum* (Schuster et al., 2020), in Arabidopsis, *rbcL* translation is increased after a shift to high light in young plants (Figure S1, Figure 3G) and in young leaves of 7-week-old plants (Figure 3H, Figure 4). However, using DMS-probing of high light-treated young plants, we observed a slight decrease of DMS reactivity at the Shine-Dalgarno sequence and no structural change at the start codon of *rbcL* (Figure 3A,C,E). Furthermore, in high light-treated young leaves, the Shine-Dalgarno sequence and the start codon show a reduction in SHAPE reactivity, indicating that the translation initiation region of *rbcL* is more compactly folded and less accessible in high light conditions (Figure 3B,C,E). In this case, our data does not support that translation initiation is regulated by the accessibility of the Shine-Dalgarno sequence and start codon. Moreover, we also did not observe significant changes in the accessibility of the sequences that have the potential to interact with the start codon and the Shine-Dalgarno sequence (Figure 3D,F).

**Figure 3.**
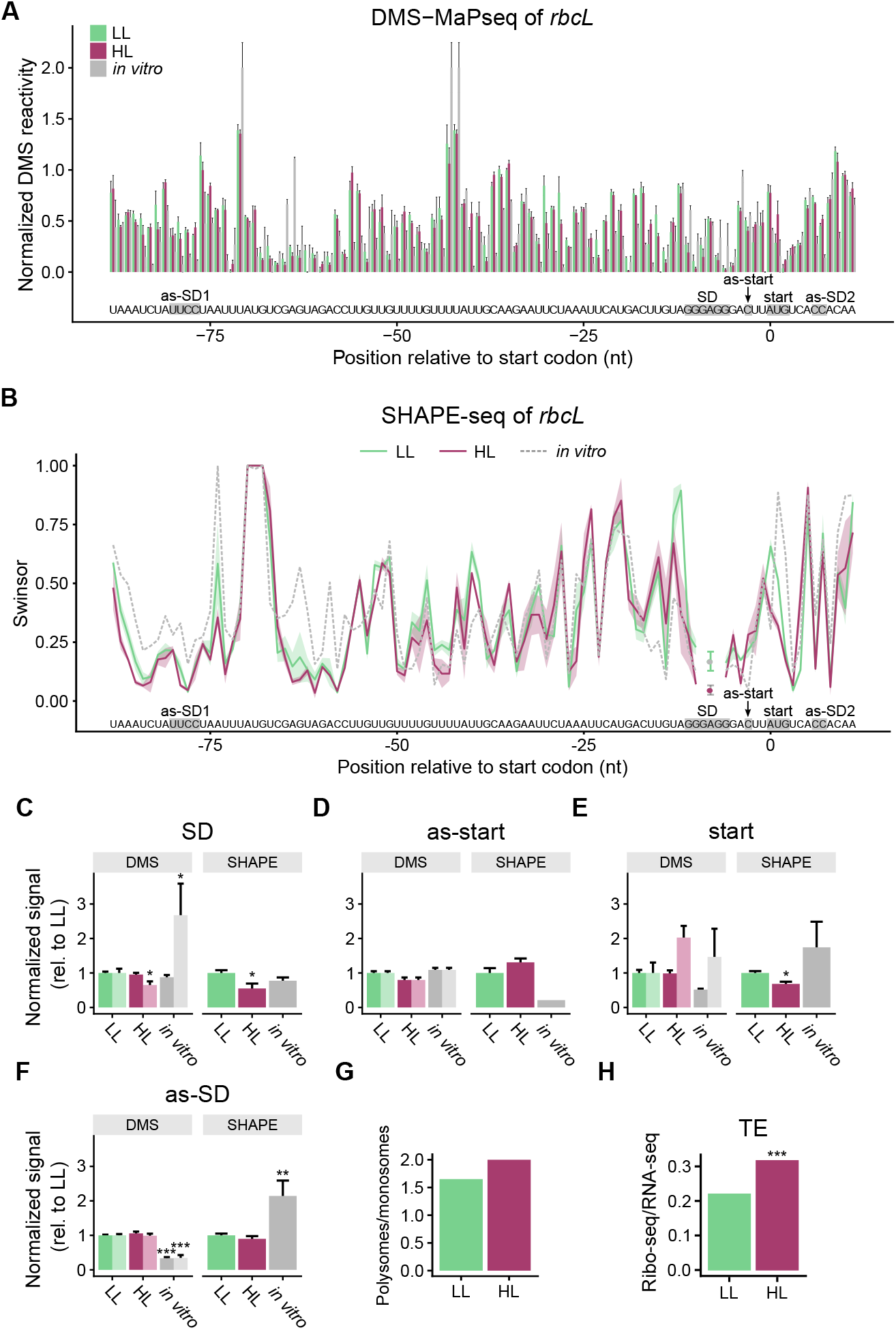
mRNA secondary structure changes in the translation initiation region of *rbcL*: **A**, DMS probing of RNA from young, 17-18-days-old plants exposed to low light (LL, light green), high light (HL, dark red), and of *in vitro*-folded RNA (gray). Normalized DMS reactivities are shown (compare Figure 1). The error bars indicate the mean standard error. The start codon (start), Shine-Dalgarno sequence (SD), and sequences that can bind the start codon (as-start) and the SD (as-SD1 and as-SD2) are marked. One portion of the SD can bind to a sequence in the coding region (as-SD2), the other one to a sequence upstream in the 5’ UTR (as-SD1). A comparison of the DMS-probed RNA with a water-treated control is shown in Supplemental Figure S4B. **B**, SHAPE analysis (NAI-N_3_ probing) of RNA from young leaf tissue obtained from 7-week-old plants presented as swinsor normalized termination counts. The shaded areas around the lines indicate the mean standard error. Positions −9 and −7 were not analyzed, because for at least one of the LL or HL samples the swinsor value was missing. **C**, reactivities at adenosines/cytidines, the columns in lighter colors are the reactivities at all four nucleotides. Asterisks here (and in **D** to **F**) indicate statistically significant changes compared to LL (*P*-values calculated with the Wilcoxon rank sum test; * = *P* < 0.05, ** = *P* < 0.01, and *** = *P* < 0.001), error bars indicate mean standard error. **D**, Average mRNA secondary structure at the nucleotide binding the start codon (as-start). **E**, Average structure at the start codon (start). **F**, Average structure at the sequences binding the SD (as-SD1 and as-SD2). **G**, Change in translation in 17-18-days-old plants (polysome fractions/monosome fractions, Figure S1). **H**, Change in translation efficiency (ratio footprints/transcript levels, Figure 4) in young leaves of 7-week-old plants. Asterisks indicate statistically significant changes compared to LL (calculated with RiboDiff; *** = *P* < 0.001).

**Figure 4.**
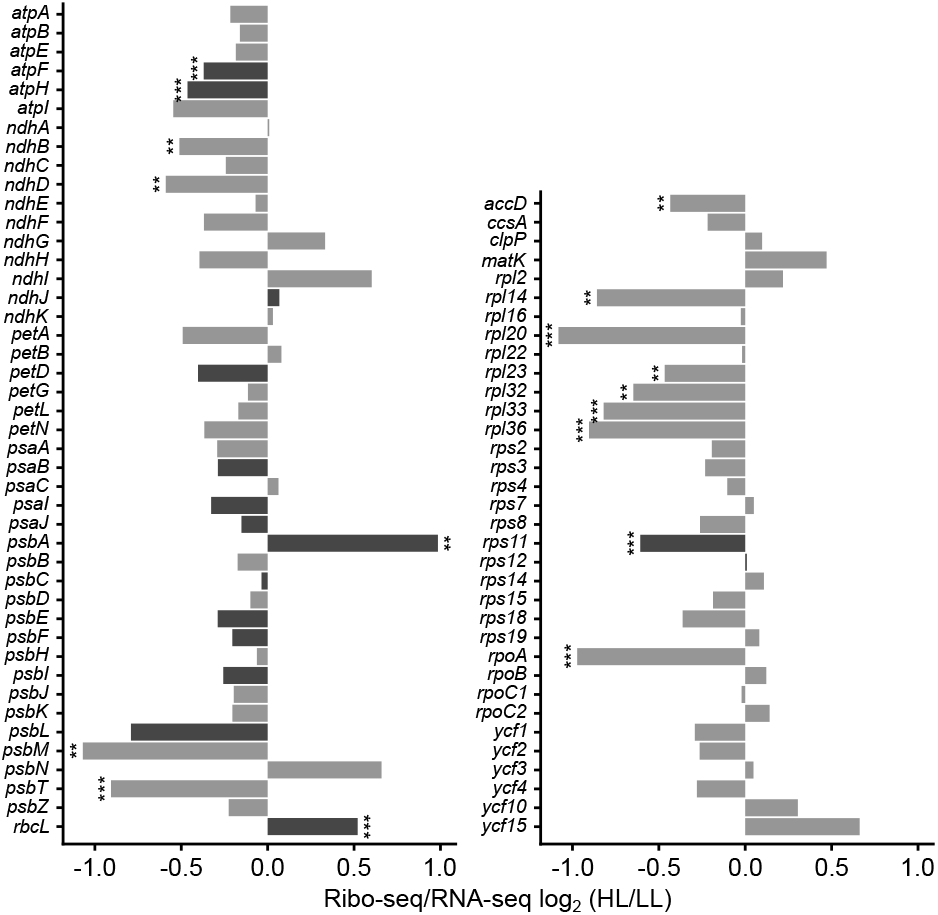
Changes in translation efficiency in response to high light treatment: The translation efficiencies (Ribo-seq/RNA-seq) for all plastid-encoded genes are shown as the ratios of the high light (HL) to the low light (LL) scores, expressed as log_2_ values. Young leaves of 7-week-old plants were analyzed (as in Figure 1C and 3B). The left panel lists the genes coding for subunits of the photosynthetic complexes, the right panel shows the data for all other genes. Translation efficiency was determined from normalized read counts for the ribosomal footprints divided by those for the transcripts of each coding region (Figure S13). Asterisks indicate statistically significant changes (calculated with RiboDiff; ** = *P* < 0.01 and *** = *P* < 0.001). Genes with bars in darker gray had sufficient coverage to permit the analysis of mRNA secondary structure (Figures 1, 3, 5).

**Figure 5.**
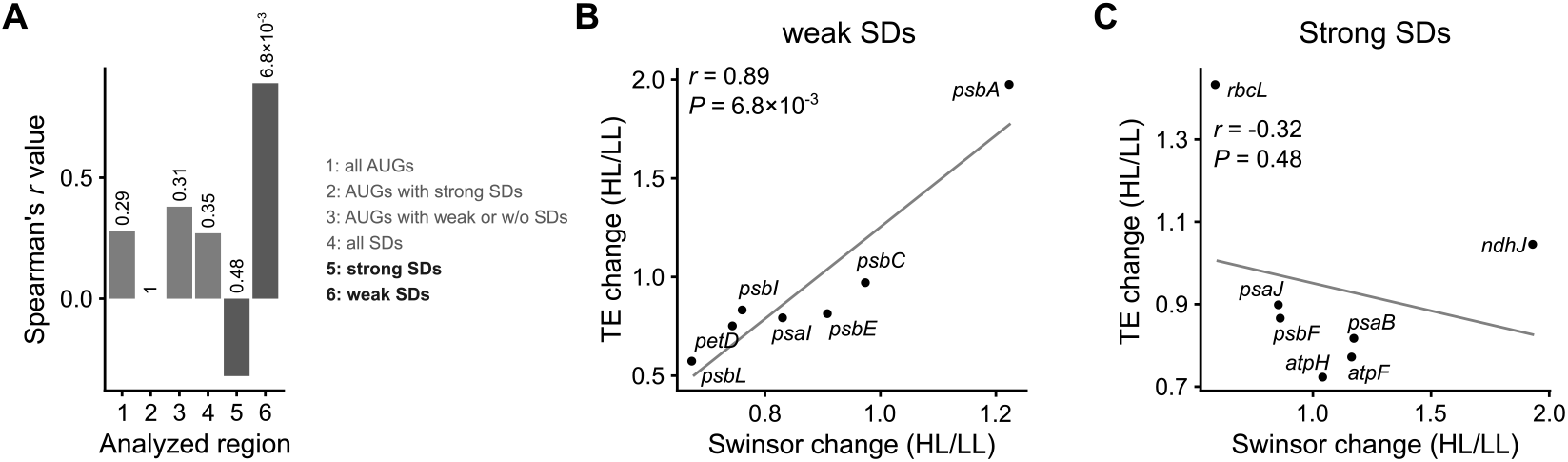
Correlations between changes in mRNA secondary structure and translation efficiency: The changes in mRNA secondary structure are calculated from the swinsor-normalized termination count values derived from NAI-N_3_ probing by dividing the values from the high light exposed plants by those from the low light control plants (young leaves of 7-week-old plants). An increase of the swinsor value indicates a decrease in base pairing, i.e. less RNA secondary structure. Average changes for the indicated segments of each gene are given. The change in translation efficiency is calculated by dividing the normalized read counts for the ribosomal footprints by those for the transcripts of each coding region, and then dividing the resulting values from the high light treatment by those from the low light control. Only genes with sufficient coverage of the mRNA secondary structure (on average at least 10 reverse transcription stops per nucleotide) are included. Spearman’s *r* and *P* values are given. **A**, Overview including all analyzed correlations. Columns 1-6 show Spearman’s *r* for the correlation between the change in translation efficiency and the change in SHAPE reactivities for different gene regions. The corresponding *P* values are given above the respective column. (1) start codons (AUG); (2) start codons of genes with strong Shine-Dalgarno sequences (SDs) (hybridization to the anti-SD of the 16S rRNA < −9 kcal mol^−1^); (3) start codons of genes with weak or no SD (> −6 kcal mol^−1^); (4) SDs; (5) SDs of genes with strong SD (< −9 kcal mol^−1^); and (6) SDs of genes with weak SD (> −6 kcal mol^−1^). The plots for all these analyses can be found in Supplemental Figure S14, where also an analysis of additional regions is included. The plots for the highlighted correlations (5) and (6) are shown in **B** (change of structure at the SDs of genes with strong SD) and **C** (SDs of genes with weak SD).

### mRNA secondary structure and translation efficiency

As shown above, structural changes of mRNA seem to be important for the high light induced translational activation of the *psbA* mRNA but not the *rbcL* mRNA (Figure 1, 3). We therefore wanted to see if there is a general correlation between structural changes and translation efficiency, or if this is a phenomenon unique to the *psbA* mRNA. Our SHAPE probing experiment of young leaves of 7-week-old plants had sufficient sequencing coverage to allow the analysis of 16 genes, including *psbA* and *rbcL*. Using the same plant material, the translation of these genes in the same plants was analyzed by ribosome profiling. This method is based on the sequencing of nuclease-protected mRNA footprints of ribosomes, which provide, when quantified per reading frame, a proxy for the synthesis rate of the corresponding protein (Ingolia et al., 2009). The reproducibility between replicates was good (Supplemental Figure S12). The translation efficiency was calculated by dividing the amount of ribosome footprints for each reading frame by the transcript levels determined by RNA-seq (Supplemental Figure S13). For several genes, a statistically significant reduction in translation efficiency was noted in high light (Figure 4). An exception was *psbA* whose translation efficiency was increased (Figure 4), which indicates increased translation initiation (Chotewutmontri and Barkan, 2018; Schuster et al., 2020) and is in accordance with the results of our polysome analysis (Supplemental Figure S7).

If a large proportion of mRNAs is regulated through RNA structural changes similar to what we observed for *psbA* upon exposure to high light, a correlation would be expected between the changes in translation efficiency and the structural alterations at the start codon and/or Shine-Dalgarno sequence (SD). However, this is not the case for either the start codons (Figure 5A-1; Supplemental Figure S14A) or the SDs (Figure 5A-4; Supplemental Figure S14D). *psbA* and *rbcL* show a higher translation efficiency in light; yet, a clear correlation with mRNA structure changes can only be found for *psbA* (Figure 1, 3). Interestingly, these two genes differ strongly regarding the strength of their SD: *rbcL* possesses a strong SD (hybridization to the anti-SD of the 16S rRNA −12.98 kcal mol^−1^), whereas the SD of *psbA* is much weaker (−5.50 kcal mol^−1^) (Supplemental Table 2). Regarding the strength of their SD, the 16 genes analyzed can be separated into a group with strongly interacting SDs (hybridization to the anti-SD of the 16S rRNA < −9 kcal mol^−1^) and a group with weak or no SDs (> −6 kcal mol^−1^) (Supplemental Table 2). In our set of 16 genes, there were only two genes without an SD, *rps11* and *rps12* (coding for the ribosomal proteins uS11c and uS12c, respectively). Therefore, SD-independent translation could not be investigated specifically, and these two genes were included in the group with weak or no SDs as appropriate. Using these two groups of genes for an analysis of the start codons, we still did not observe a significant correlation between the changes in SHAPE reactivities and the change in translation efficiency (Figure 5A-2,A-3; Supplemental Figure S14B,C). In contrast, there was a clear difference between the groups regarding the structure at the SD. Genes with weak SDs showed a statistically significant correlation between the change in translation efficiency and the change in SHAPE reactivities in the SDs (Figure 5A-6, 5B; Supplemental Figure S14F). No such correlation was observed for genes with strong SDs (Figure 5A-5, 5C; Supplemental Figure S14E).

Thus, our data (Figure 1, 2, 5B) suggest that the structural accessibility of the SD region is central for the light-dependent translational regulation of mRNAs with weak SDs (such as *psbA*), whereas other mechanisms are likely to be more important for mRNAs with strong SDs (Figure 3, 5C). In the case of the *psbA* mRNA, translational regulation seems to depend on the recruitment of specific proteins to the 5’ UTR region and subsequent remodeling of the RNA structure.

## DISCUSSION

The molecular mechanisms of translation regulation in plastids of higher plants have been elusive. *In vitro* data showed that binding of putative regulatory proteins influences the mRNA secondary structure of the region encompassing the start codon and/or the Shine-Dalgarno sequences (SD) (Prikryl et al., 2011; Hammani et al., 2012). It was postulated that such a mechanism might act to regulate translation *in vivo*. We tested this hypothesis by analyzing the secondary structure and translation efficiency of plastid mRNAs from plants exposed to low and high levels of light.

In high light, *psbA* was the plastid mRNA with the strongest increase in translation efficiency (Figure 4) – as expected because the turnover of its protein product, D1, increases under these conditions (Li et al., 2018). Regulation of *psbA* translation differs between dark/light shifts and the response to increasing D1 turnover (PSII repair) (Zoschke and Bock, 2018; Chotewutmontri and Barkan, 2018). At least in higher plants, the regulation in response to dark/light shifts happens on the level of translation elongation (Chotewutmontri and Barkan, 2018), whereas under conditions of high D1 turnover, *psbA* translation is induced on the level of translation initiation (Chotewutmontri and Barkan, 2018; Schuster et al., 2020), as indicated also by polysome analysis (Supplemental Figures S1, S7) and ribosome profiling (Figure 4). The *cis* elements required for initiation of *psbA* translation are not strongly conserved in higher plants: The *psbA* mRNA in Arabidopsis has a weak SD, whereas in some other species, e.g. *Nicotina tabacum* and *Zea mays*, *psbA* completely lacks a SD (Supplemental Table 2) (Scharff et al., 2017). In contrast, the *trans* factors regulation psbA translation are probably conserved in higher plants: Three proteins have been reported to activate *psbA* translation: HCF173 (Schult et al., 2007), HCF244 (Link et al., 2012), and LPE1 (Jin et al., 2018), whereas AtPDI6 is described as a negative regulator (Wittenberg et al., 2014). Furthermore, also the chlorophyll-binding proteins OHP1 and OHP2 are important for translation activation of *psbA* (Chotewutmontri et al., 2020). However, conflicting results indicate that LPE1 binds to *psbJ* and *psbN*, not *psbA* (Williams-Carrier et al., 2019), and LPE1 was not found to be bound to *psbA* mRNA (McDermott et al., 2019). HCF173 was described to be one of the proteins contributing to the footprint detected in the *psbA* 5’ UTR (Figure 2, Supplemental Figure S11A,D) (McDermott et al., 2019). D1 is inserted co-translationally into thylakoid membranes. HCF173 and HCF244 are bound to the thylakoids (Link et al., 2012). HCF244 is possibly recruited there via an interaction with OHP1 and OHP2 (Hey and Grimm, 2018; Myouga et al., 2018). If HCF173, HCF244, LPE1, AtPDI6, OHP1 and/or OHP2 are involved in the regulation of *psbA* translation, it could be assumed that these proteins themselves and/or their expression are subject to light-dependent regulation. However, we did not observe any alterations in the transcript levels and translation efficiency of their genes during high light acclimation (Supplemental Table 3), indicating that light-dependent regulation of these proteins, if it occurs, must take place post-translationally. How the described proteins might activate *psbA* translation, either alone or as a complex, was unknown.

Using DMS and a SHAPE reagent, NAI-N_3_, we demonstrated that the degree of secondary structure of the Shine-Dalgarno sequence and the start codon in the *psbA* mRNA is reduced *in vivo* under high light conditions and that this correlates with increased translational efficiency (Figure 1). This correlation is compatible with the hypothesis that translation is activated by making the SD and/or start codon more accessible. Furthermore, we and others found evidence for a possible binding site for a regulatory protein in a position where binding could result in structural changes of the translation initiation region as predicted by the hypothesis (Figure 2) (McDermott et al., 2019). These findings argue that the regulation of *psbA* translation could involve the modulation of mRNA secondary structure by protein binding.

There are indications that such a mechanism is used by other genes: in the case of genes with weak SDs, the change in mRNA secondary structure at the SD correlates with the change in translation efficiency (Figure 5B). Interestingly, the correlation is specific for genes with weak SDs. It is possible that strong SDs are more likely to hybridize to the anti-SD of the 16S rRNA, and therefore are less amendable to regulation by alternative mRNA secondary structures. Accordingly, *rbcL* is an example of a gene with a strong SD (Supplemental Table 2) and here the increased translation efficiency under high light conditions cannot be explained by changes in the structure of the SD and the start codon (Figure 3). Furthermore, the comparison of the structural changes in the translation initiation regions of *psbA* (Figure 1) and *rbcL* (Figure 3) indicates that the structure alterations are not a consequence of increased translation itself, e.g. by increased binding of the ribosome (including tRNA-fMet(CAU)) at the start codon. Both genes are upregulated at the level of translation, but the degree of secondary structure does not change in the same direction. Therefore, these structural changes also cannot be caused simply by increased temperatures during high light treatment; in the case of *rbcL,* the SD and start codon are not paired to a lower extent in high light, whereas heat would normally be expected to decrease pairing. How translation of *rbcL* itself is regulated remains unknown. It is possible that distinct mechanisms for regulation of translation initiation exist as plastids use two distinct mechanisms for start codon recognition (Scharff et al., 2011, 2017).

The results for *psbA* and other plastid genes with weak SDs are in agreement with reports for *E. coli* that translation efficiency is determined by the extent of RNA secondary structure at the SD (Bhattacharyya et al., 2018; Mustoe et al., 2018). In bacteria, several mechanisms are described to regulate translation initiation by altering the accessibility of SDs, including RNA thermometers (Neupert et al., 2008; Krajewski and Narberhaus, 2014), binding of small RNAs and proteins (Laursen and Sørensen, 2005; Duval et al., 2015), and riboswitches (Breaker, 2018). Synthetic riboswitches are also functional in plastids (Verhounig et al., 2010). Our results (Figure 1, 5B) suggest that in plastids a similar mechanism, based on the manipulation of mRNA secondary structure by RNA-binding proteins, is used for the regulation of translation of *psbA* and other genes with weak SDs.

## EXPERIMENTAL PROCEDURES

### Plant material

For DMS probing, *Arabidopsis thaliana* wild-type (ecotype Col-0) plants were grown in Jiffy pots (Jiffy Products) for 17-18 days at 22 °C and 150 μE m^−2^ s^−1^ in long-day conditions (16 h day/8 h night). Then they were either kept for 1 h in dim light (~10 μE m^−2^ s^−1^, low light control) or shifted for the same time to 1000 μE m^−2^ s^−1^ white light (high light treatment) supplied by an SL 3500-W-D LED lamp (Photon Systems Instruments). Plants treated in the same way were used for polysome analysis (Supplemental Figure S1).

For NAI-N_3_ probing, *A. thaliana* plants were grown for 7 weeks at 20 °C in short-day (8 h day/16 h night), low light conditions (140-160 μE m^−2^ s^−1^). The low light sample was harvested at noon, while the high light sample was transferred at noon to the following conditions: 4 h high light [1200 μE m^−2^ s^−1^]; 16 h dark; 4 h high light [1200 μE m^−2^ s^−1^]; and then the leaf material was harvested. The temperature of the growing chamber was set to 20 °C, but owing to the heat emitted by the lamps, the leaves were exposed to temperatures of up to 30 °C. Young leaves with a maximum length of 20 mm were harvested into liquid nitrogen (rosette diameter at this growth stage was 68 ± 3 mm). Plants treated the same way were used for ribosome profiling and RNA-seq (Figure 1G, 3H, 4, Supplemental Figure S11A-D) as well as polysome analysis (Supplemental Figure S7).

### Determining photosynthetic parameters

Chlorophyll *a* fluorescence parameters were measured in triplicates using a MAXI IMAGING-PAM M-series instrument (Walz). Plants were dark-acclimated for 30 min. For F_0_ and F_m_ determination, plants were exposed to a saturating pulse followed by 5 min of blue (450 nm) actinic light (81 μE m^−2^ s^−1^). In an actinic light phase, saturating light pulses were applied at 20-s intervals. Results were calculated for the last saturating pulse during the actinic light period. Maximum quantum yield of photosystem II (F_v_/F_m_) and electron transport rate (ETR) parameters were calculated as described previously (Klughammer and Schreiber, 2008).

### Polysome analysis

Polysome analysis using sucrose gradients for separation of free mRNA and polysome complexes was done as described previously (Barkan, 1993). The *psbA* and *rbcL* probes were amplified from total plant DNA using gene-specific primers (see Supplemental Table 4), radioactively labelled with α32P[CTP] using the Megaprime DNA Labeling System (GE Healthcare Life Sciences), and hybridized at 65 °C.

### Ribosome profiling (Ribo-seq)

Ribosome profiling was done as described before (Oh et al., 2011; Zoschke et al., 2013; Gawroński et al., 2018). Three biological replicates (each consisting of material from at least three plants) for each treatment were analyzed as follows: 400 mg of deep-frozen, ground leaf material was thawed on ice in 5 ml of extraction buffer (200 mM Tris/HCl pH 8.0, 200 mM KCl, 35 mM MgCl_2_, 0.2 M sucrose, 1% Triton X-100, 2% polyoxyethylen-10-tridecyl-ether, 5 mM dithiothreitol, 100 μg/ml chloramphenicol, 50 μg/ml cycloheximide). The extract was centrifuged for 5 min at 13,200 g and 4 °C. 600 μl of the supernatant was removed for analysis by RNA-seq (see below), and the remaining supernatant was centrifuged for 10 min at 15,000 g and 4 °C. CaCl_2_ was added to the resulting supernatant to a concentration of 5 mM, followed by 750 units of micrococcal nuclease (Thermo Fisher), and the mixture was incubated for 1 h at room temperature. The digested extract was loaded on a 2-ml sucrose cushion (40 mM Tris/acetate pH 8.0, 100 mM KCl, 15 mM MgCl_2_, 1 M sucrose, dithiothreitol, 100 μg/ml chloramphenicol, 50 μg/ml cycloheximide) and centrifuged for 3 h at 55,000 g and 4 °C in a Type 70 Ti rotor (Beckman). The pellet was dissolved in 1% SDS, 10 mM Tris/HCl pH 8.0, and 1 mM EDTA. RNA was purified using the PureLink miRNA Isolation Kit (Invitrogen). The 16 to 42-nt fraction was isolated by electrophoresis and treated with T4 polynucleotide kinase before library preparation using the TruSeq Small RNA Library Preparation Kit (Illumina). Sequencing was performed on the HiSeq 4000 platform (Illumina).

### RNA-seq

For each treatment three biological replicates were analyzed. RNA was purified from 600 μl of leaf extract (see ribosome profiling) using the RNAeasy Plant Mini Kit (Qiagen). The RNA was treated with the Ribo-Zero™ rRNA Removal Kit (Plant Leaf) (Illumina), libraries were prepared using the TruSeq RNA Library Prep Kit v2 (Illumina) and sequenced on the HiSeq 4000 platform (Illumina).

### Determination of 3’-ends of plastid transcripts

Determination of 3’-ends was done with the protocol described previously (Marquardt et al., 2014). Briefly, a DNA linker (NEB) was ligated to free 3’-ends of 1 μg of denatured total RNA. After ligation, the RNA was fragmented in an alkaline solution (100 mM NaCO_3_, 2 mM EDTA) for 30 min. The RNA was subsequently precipitated, dissolved, and loaded on a 15% TBE-Urea gel. Fragments in the size range of 50-150 bp were cut out and precipitated overnight. The fragmented RNA was then used to synthesize cDNA with Superscript III (Invitrogen) and a primer that annealed to the ligated linker. The cDNA was loaded on a 10% TBE-Urea gel and products in the size range 85-160 bp were cut out and precipitated. Products were then circularized with CircLigase (Epicentre) according to the manufacturer’s instructions and used as a template for PCR amplification. The reaction included a primer with a barcode for sequencing purposes. Amplified PCR products were loaded on an 8% TBE gel and products around 150 bp were cut out. The sequencing library was run on a Bioanalyzer (Agilent) to confirm that no contaminations from the library construction were present. Sequencing was done on a MiSeq platform (Illumina).

### Processing of Ribo-seq and RNA-seq reads

*Arabidopsis thaliana* (Col-0) genomic, transcriptomic and non-coding RNA sequences, and the GFF3 annotation file were downloaded from Ensembl Plants (http://plants.ensembl.org, release 41). This annotation file lacks the annotation of the plastid transcripts. We added our own annotation using a manually curated data set. The 5’ ends are based on primer extension data from the RNA secondary structure probing with NAI-N_3_ (see below). The 3’ ends are based on the 3’-end mapping set (see above). If there are multiple transcripts for one gene, the longest transcript detected was chosen for the annotation file. The sequences of coding regions were corrected for editing as detected by RNA-seq. Start codons and missing exons were corrected using GeSeq (Tillich et al., 2017) plus corrections based on the ribosome profiling data. *rps16* was not spliced and was therefore characterized as a pseudogene as described previously (Roy et al., 2010). From the downloaded transcriptome, plastid sequences were replaced with the new set.

Adapter sequences were removed using TrimGalore! (version 0.4.5; http://www.bioinformatics.babraham.ac.uk/projects/trim_galore/). Alignments were performed with STAR (version 2.6.0a;) (Dobin et al., 2013) with following settings: `--outFilterMismatchNmax 2 --outMultimapperOrder Random --outSAMmultNmax 1 --alignIntronMax 1 – alignIntronMin 2` allowing for two mismatches, ungapped alignment on the transcriptome, and random assignment of reads that mapped to more than one location. Reads that mapped to non-coding RNAs were removed from the analysis. Unaligned reads were used as an input in an alignment to the transcriptome. Reads whose alignment length was between 28 nt and 40 nt and which mapped in a “sense” direction were used for further analysis. To assign each footprint to the P-site of the ribosome, we used the 5’-end of a mapped footprint and 23-nt offset as described previously (Gawroński et al., 2018). Only reads whose P-site overlapped with CDSs were used for further analysis. When a P-site overlapped with more than one CDS (e.g. in partially overlapping *psbD*-*psbC*), the read was assigned randomly to one of the CDSs. RNA-seq reads were mapped to the transcriptome in a similar way, but reads with more than 5% mismatches were removed. Reads that mapped in both directions (unstranded library) and those that overlapped by at least 1 nt with CDSs were used for further analysis. Similarly to Ribo-seq, random assignment was used when a read overlapped with more than one CDS (e.g. *psbD*-*psbC*). Based on counts of reads mapped to the CDSs, RPKM (reads per kilobase per million mapped reads) values were calculated using normalization to the total number of mapped reads for each sample and the length of the CDS. For the analysis of footprints of putative regulatory proteins, reads with an aligned length of between 18 nt and 40 nt were used.

### Calculation of translation efficiency and analysis of differential gene expression

Translation efficiency (TE) was calculated using RiboDiff (version 0.2.1) (Zhong et al., 2017) and counts of reads were mapped to the CDSs. Genes with *P* < 0.01 were considered to be significantly changed. For the differential gene expression analysis, RNA-seq reads were pseudo-aligned to the transcriptome using Salmon (version 0.9.1) (Patro et al., 2017) with default parameters. Transcript-level abundances were imported into R using tximport (Soneson et al., 2015) and analyzed using the DESeq2 package (Love et al., 2014).

### Gel-blot analysis of small RNAs

RNA was extracted from leaf material harvested in low light and high light (same material as used for ribosome profiling, RNA-seq, and RNA secondary structure probing with NAI-N_3_) by adding 666 μl of extraction buffer (see ribosome profiling) to frozen, ground material. The RNA was purified from the extract using a phenol/chloroform/isoamyl alcohol step and isopropanol precipitation. The gel blot was done as described before (Loizeau et al., 2014): 10 μg total RNA was separated on 15% polyacrylamide TBE urea gels (Biorad) and transferred in a wet-blot setup with 0.5 TBE buffer to a Hybond-N membrane (GE-Healthcare). The RNA was cross-linked to the membrane with 0.16 M N-(3-Dimethylaminopropyl)-N′-ethylcarbodiimide hydrochloride in 0.13 M 1-methylimidazole (pH 8.0) at 60 °C for 1 h. The probe (see Supplemental Table 4) was labelled at the 5’ end with γ-^32^P-ATP and hybridized at 60 °C using standard protocols.

### RNA structure probing with DMS

Three biological replicates (each consisting of at least three plants) for each treatment were used. Young, 17-18 days old plants were collected into 10 ml of DMS reaction buffer (100 mM KCl, 40 mM HEPES pH 7.5, 0.5 mM MgCl_2_). Dimethyl sulfate (DMS) was added to a concentration of 5% (w/v) and the reaction was performed at 24-25 °C (DMS+ samples). In parallel, negative control (DMS-) samples were prepared by adding water in place of DMS. The young plants were incubated for 6 min at either ~10 μE m^−2^ s^−1^ (low light control) or 1000 μE m^−2^ s^−1^ (high light treatment) while the solution was held horizontally and hand mixed. The high light treatment caused a 1 °C temperature increase in the reaction buffer. The reaction was stopped by adding 20 ml of ice-cold 30% β-mercaptoethanol and incubating for 1 min on ice. Afterwards the liquid was removed, and the plants were washed twice with distilled water and frozen in liquid nitrogen. RNA was extracted using the Spectrum Plant Total RNA Kit (Sigma-Aldrich). DNA was removed using the Turbo DNA-free kit (Thermo Fisher Scientific). cDNA was produced using 1-μg aliquots of RNA as template, 0.5 μM target-specific primer (see Supplemental Table 4), 100 units TGIRT-III (InGex) reverse transcriptase in TGIRT buffer (50 mM Tris-HCl pH 8.3, 75 mM KCl, 3 mM MgCl_2_), 1 mM dNTPs, 5 mM dithiothreitol, and 4 units of Murine RNase Inhibitor (NEB)). The mixture was incubated for 2 h at 57 °C. RNA was removed by adding 5 units of RNase H (NEB) and incubating for 20 min at 37 °C. RNase H was inactivated by 20-min incubation at 65 °C. cDNA was purified using 1.8X strength Ampure XP beads (Beckman Coulter). The region of interest was amplified with specific primers (see Supplemental Table 4) and the Q5 DNA polymerase (NEB), and indexed by PCR using primers containing Illumina indexes (see Supplemental Table 4), and sequenced on a MiSeq (Illumina) sequencer (2 × 300 bp).

For *in vitro* DMS probing, 5 μg (20 μl) of DNase treated RNA in water was heat denatured for 2 min at 95 °C and quickly transferred to ice. 80 μl of DMS reaction buffer (100 mM KCl, 40 mM HEPES pH 7.5, 0.5 mM MgCl2) and 100 U of Murine Rnase Inhibitor (NEB) were added, followed by incubation with mixing at 25 °C for 5 min. Next, DMS (or water for DMS- samples) was added to the final concentration of 5% and samples were incubated for 5 min at 25 °C with gentle mixing. The reaction was terminated by adding 200 μl of ice cold 30 % β-mercaptoethanol and incubating for 1 min on ice. RNA was recovered by ethanol precipitation. cDNA synthesis, library preparation and sequencing was done as described above.

### DMS-MaPseq analysis

Adapter sequences from reads were removed using TrimGalore! (version 0.4.5) with the following settings: --fastqc --quality 35 --length 75 (and -max_length 200 for reverse reads). Reads were mapped to the *psbA*, *rbcL* and 16S rRNA using bowtie2 (version 2.3.4.1) (Langmead and Salzberg, 2012) separately for forward and reverse reads with following settings: --local -very-sensitive-local -p 12 -U. Mutation frequencies for *psbA*, *rbcL*, and 16S rRNA regions located between the primers used for amplification were calculated using the pileup function from the Rsamtools package (Morgan et al., 2017). For further analysis, substitutions and deletions at nucleotides with coverage higher than 1500 reads and not bound by primers were used. Raw DMS reactivities from DMS- were subtracted from DMS+ samples and all negative values set to 0. Next, DMS reactivities were normalized separately for G/U and A/C by dividing the reactivities by mean reactivity of most highly reactive nucleotides (90th-99th percentile) of each transcript followed by 99% winsorization to remove extremely high values, as described earlier (Gawroński et al., 2020). For structure prediction, RNA sequences were folded by the Fold program from RNAstructure (version 6.2) (Mathews et al., 2016) with normalized DMS reactivities for all nucleotides used as soft constrains. Fold program parameters were as follows: -md 500 -t 298.15. For the *psbA* high light samples, the protein binding site was forced to be single-stranded. Structures were visualized using VARNA (Darty et al., 2009).

### Structure analysis of 16S rRNA

The crystal structure of the chloroplast 70S ribosome (Ahmed et al., 2017) was downloaded from PDB (https://www.rcsb.org/, entry 5X8P). Surface residues (i.e. solvent accessible) were calculated in PyMOL using the FindSurfaceResidues module. Residues with an area > 2.5 Å^2^ were considered as solvent accessible.

### RNA secondary structure probing with NAI-N_3_

Ten samples in all were prepared, eight of which were derived from plant material grown under low light control conditions and two from high light material. All but one of the samples were structure-probed using the SHAPE reagent NAI-N_3_; the exception was exposed to a mock treatment using DMSO as a probing control (Flynn et al., 2016). All samples, except the DMSO control and one low light sample, were selected for probing of induced termination (Poulsen et al., 2015). One low light sample was subjected to probing of *in-vitro*-folded RNA. The others were probed using homogenized, flash-frozen leaf tissue, imitating *in-vivo* conditions.

The sample probed *in vitro*, as well as two low light and the two high light samples, were depleted of rRNA, whereas all others were comprised of total RNA. For low light conditions, there were two biological replicates each for the total RNA and the rRNA-depleted RNA. For the total RNA, an additional technical replicate was generated by splitting the sample after DNase treatment (see below). For the high light conditions, two biological replicates were analyzed.

A 2 M NAI-N_3_ stock solution was prepared by mixing dropwise 0.15 g of 2-azidomethyl nicotinic acid dissolved in 210 μL DMSO with 0.14 g of carbonyldiimidazole in 210 μL DMSO, and letting the two react for 1 h. Probing was done by adding to 100 mg of deep frozen, ground leaf material to 540 μl extraction buffer (0.92 M HEPES/KOH pH 8.0, 5 mM MgCl_2_, 0.5 mg/ml heparin, 1% Triton X-100, 2% polyoxyethylen-10-tridecyl-ether) pre-mixed with 60 μl of 1 M NAI-N_3_ in DMSO (giving a final concentration of 100 mM). The sample was incubated for 2 min at room temperature. The reaction was stopped by addition of β-mercaptoethanol to a final concentration of 1.4 M. Cell debris was removed by centrifugation for 5 min at 13,200 g and 4 °C. RNA was isolated using phenol/chloroform/isoamylalcohol (25:24:1) extraction and isopropanol precipitation. DNA was removed using the RNase-Free DNase Set (Qiagen) according to the manufacturer’s protocol. For some of the samples (see above), rRNA was depleted using the Ribo-Zero Bacterial rRNA Removal Kit (Illumina). To evaluate the efficiency of subsequent selection of probed RNA, before cDNA synthesis 1% and 2% of *E. coli fhlA220* mRNA was spiked into total RNA and rRNA-depleted samples, respectively. The cDNA synthesis was carried out with modifications as described (Takahashi et al., 2012). Specifically, 1 μl 50 μM random primer (RT_15xN, see Supplemental Table 4) was annealed to 8 μL of total RNA or rRNA-depleted RNA by incubation at 65 °C for 5 min and then transferred to ice. A 28-μL aliquot of a mastermix consisting of transcription buffer (250 mM HEPES pH 8.3, 375 mM KCl, 15 mM MgCl_2_), 7.5 μl 2.5 mM dNTPs, 7.5 μl sorbitol (3.3 M)/trehalose (0.6 M), 500 units PrimeScript Reverse Transcriptase (Takara), and 3 μl water was added to each sample. Samples were then incubated at 25°C for 30 s, 42°C for 30 min, 50°C for 10 min, 56°C for 10 min, and 60°C for 10 min, and subsequently purified using XP RNA beads (Ampure) according to the manufacturer’s protocol. The samples were biotinylated as described earlier (Flynn et al., 2016). Full-length cDNA was selected using 100 μl MPG Streptavidin beads (PureBiotech) per sample as described (Takahashi et al., 2012) with minor alterations. The beads were blocked with 1.5 μl of a 20 μg/μl *E. coli* tRNA mix for 60 min at room temperature, separated from the supernatant on a magnetic stand, and washed twice with 50 μl of wash buffer 1 (4.5 M NaCl, 50 mM EDTA pH 8.0), followed by resuspension in 80 μl wash buffer 1. The beads were then mixed with 40 μl of cDNA/RNA sample and incubated at room temperature for 30 min with vortexing every 10 min. After 5 min on a magnetic stand, the supernatant was removed, and the beads were washed a total of 6 times with 150 μL of the following wash buffer: 1x wash buffer 1, 1x wash buffer 2 (300 mM NaCl, 1 mM EDTA pH 8.0), 2x wash buffer 3 (20 mM Tris-HCl pH 8.5, 1 mM EDTA pH 8.0, 500 mM NaOAc pH 6.1, 0.4% SDS), 2x wash buffer 4 (10 mM Tris-HCl pH 8.5, 1 mM EDTA pH 8.0, 500 mM NaOAc pH 6.1). To release the cDNA from the beads, 60 μl of 50 mM NaOH was added and the samples were incubated for 10 min at room temperature. The eluate was removed after separation on a magnetic stand and mixed with 12 μL of 1 M Tris-HCl (pH 7), followed by ethanol precipitation. Libraries were prepared as previously described (Poulsen et al., 2015) with minor modifications. A mixture consisting of 1 μl 10x Circligase reaction buffer (Epicentre), 0.5 μl 1 mM ATP, 0.5 μl 50 mM MnCl_2_, 2 μl 50% PEG 6000, 2 μl 5 M betaine, 0.5 μl 100 μM Ligation_adapter oligonucleotide (see Supplemental Table 4), and 50 units of Circligase (Epicentre) was added to 3 μl of cDNA and incubated at 60°C for 2 h and 68°C for 1 h, followed by enzyme inactivation at 80°C for 10 mins. The ligated cDNA was purified by ethanol precipitation and resuspended in 20 μl H_2_O, of which 5 μl was used for PCR with 45 μl PCR reaction mix (3 μl of PCR_forward primer, 2.5 μl of indexed reverse primer (see Supplemental Table 4), 10 μl of Phusion 5× HF buffer, 4 μl of 2.5 mM dNTPs, 24.5 volume H_2_O, and 2 units Phusion Polymerase (NEB)). The PCR was conducted with the following cycles: 1× (98°C for 3 min), 5× (98°C for 80 s; 64°C for 15 s; 72°C for 1 min), 16× (98°C for 80 s; 72°C for 45 s), 1× (72°C for 5 min) and purified using Ampure XP beads, eluting the PCR product in 30 μl water. The molar distribution of the individual samples was analyzed using a Bioanalyzer High sensitivity chip (Agilent) and used to pool samples equally followed by size selection (200–600 bp range) on an E-gel 2% SizeSelect gel (Invitrogen). The size-selected library was precipitated and resuspended in 20 μl of water followed by Ampure XP bead (ratio 1:1.8) purification. The library was sequenced on the Illumina NextSeq system with the 75 bp single-end protocol.

### In vitro RNA secondary structure probing with NAI-N_3_

DNA-depleted RNA was folded *in vitro* and SHAPE probed as described (Flynn et al., 2016), with modifications. Specifically, 10 μg RNA in water was heat-denatured for 2 min at 95 °C and transferred to ice. SHAPE reaction buffer (100 mM HEPES pH 8.0, 6 mM MgCl_2_, 100 mM NaCl) and 400 units of RiboLock RNAse inhibitor (Thermo Fisher Scientific) were then added, followed by incubation for 5 min at 37 °C. Subsequently, NAI-N_3_ was added to a final concentration of 100 mM, followed by gentle mixing and incubation at 37 °C for 10 min. The reaction was terminated with β-mercaptoethanol (1.4 M final concentration) and the RNA was recovered by ethanol precipitation. Reverse transcription, biotinylation, selection of probed sequences, library preparation, and sequencing were done as described above.

### SHAPE data analysis

Data analysis was conducted either on a debian Linux server as command line functions or in RStudio (V. 1.1.456). Adapter sequences, short reads, and low quality 3’-ends were removed from the reads using cutadapt v. 1.15 (Martin, 2011) with the options `cutadapt -a GATCGGAAGAGCACACGTCT --quality-cutoff 17 --minimum-length 40`. The random barcodes incorporated into the 3’ adapter were removed and saved for later analysis using `preprocessing.sh` (Kielpinski et al., 2015) with the options `-b NNNNNNN` and `-t 15` for barcode and trimming length respectively. Sequenced reads were mapped using Bowtie2 v2.2.3 (Langmead and Salzberg, 2012) with the options `--norc -N 1 -D 20 - R 3 -L 15`. The reads were mapped to a fasta file containing manually annotated Arabidopsis transcripts (see Ribo-seq and RNA-seq reads processing). Using the barcode and sequence information, the counts from observed unique barcodes were summarized with `summarize_unique_barcodes.sh` with the options `-t -k` to trim untemplated nucleotides and to produce a `k2n` file, respectively (Kielpinski et al., 2015). To account for bias during library preparation the estimated unique barcodes were calculated with the R package “RNAprobR” (v. 1.2.o) function readsamples() with the `euc`=“`HRF-Seq`” option (Kielpinski et al., 2015) using Rstudio Version 1.1.456. Finally, the count data from the estimated unique counts was compiled with the original fasta file to create positional information using the RNAprobR function comp(). The compiled data was subsequently normalized by a 90% winsorization, whereby all values in a sliding 51-nt window were set to the 98th percentile. Comparison of the samples revealed that two samples were extreme outliers (Supplemental Figure S8) and they were excluded from further analysis. Both were low light samples, one of which contained total RNA (LL4), the other one had been depleted of rRNA (LL5). The remaining samples included in the following analysis were: the low light samples LL1 (total RNA), LL2 (total RNA, technical replicate of LL1), and LL3 (rRNA depleted) plus the DMSO control, the not selected control and one *in-vitro*-folded sample; for high light conditions, HL1 and HL2 (both rRNA depleted). For structural analysis only genes with on average more than 10 reverse transcription stops per nt were used (Supplemental Figure S8). Positions in these genes that were missing swinsor values in at least one of the LL or HL samples were not analyzed (e.g. positions 9 and 7 in *rbcL*; Figure 3). Normalized swinsor values for selected motifs (i.e. start, SD, as-SD and as-start) were calculated by dividing, for each nucleotide, the swinsor value by the average swinsor for that nucleotide in all LL samples. The SDs were identified by *in silico* hybridization of the anti-SD CCUCCU of the 16S rRNA to nucleotides −22 to −2 of each 5’ UTR at 20 °C using Free2bind (Starmer et al., 2006). The same program was also used to determine the strength of the interaction between SDs and anti-SD, and the SDs were classified into strong and weak categories.

### Receiver operating characteristic (ROC) curve

Using the roc() function from the pROCpackage (v. 1.9.1) in R, a receiver operating characteristic (ROC) curve was generated using the dot-bracket structure from the Arabidopsis 18S rRNA obtained from the “The comparative RNA web” (CRW) site (Cannone et al., 2002) as predictor and the swinsor normalized termination counts as response. From the generated ROC curve, the area under the curve was calculated. To assess the quality of DMS data, we performed receiver operating characteristic (ROC) using pROC package (Robin et al., 2011) based on crystal structure of chloroplast ribosome (Ahmed et al., 2017), as described earlier (Gawroński et al., 2020).

### Data availability

All data analysis was performed in R (R Development Core Team, 2018) and plotted using ggplot2 (Wickham, 2016). All sequences were deposited in the Sequence Read Archive under BioProject number [will be submitted in time].

## Acknowledgements

We thank Paul Hardy and Stefanie Zintl for commenting on the manuscript. The authors acknowledge financial support from The Polish National Science Centre (Narodowe Centrum Nauki) to P.G. (SONATA12, UMO-2016/23/D/NZ3/02491) and S.K. (MAESTRO6 2014/14/A/NZ1/00218); from the European Research Council (ERC) and the Marie Curie Actions under the European Union’s Horizon 2020 research and innovation programme to S.M. (StG2017-757411) and to P.K. MSCA-IF 703085; from the Hallas-Møller Investigator award by the Novo Nordisk Foundation (NNF15OC0014202) and the Copenhagen Plant Science Centre Young Investigator Starting grant to S.M; the German Science Foundation (DFG, grant TR175) to D.L; from the Independent Research Fund Denmark (Danmarks Frie Forskningsfond; 7014-00322B) and the VILLUM Foundation (Project no. 13363) to P.E.J.

## Short legends for Supporting Information

Supplemental Figure S1 Polysome analysis of *psbA* and *rbcL* translation in young plants.

Supplemental Figure S2 RNA quality assessment after DMS treatment.

Supplemental Figure S3 DMS-MaPseq quality control including reproducibility.

Supplemental Figure S4 DMS-MaPseq compared to water control.

Supplemental Figure S5 DMS data reproduces known elements of the secondary structure of 16S rRNA.

Supplemental Figure S6 Determination of photosynthetic parameters.

Supplemental Figure S7 Polysome analysis of *psbA* translation in mature plants.

Supplemental Figure S8 Reproducibility of the mRNA secondary structure probing (NAI-N_3_) data.

Supplemental Figure S9 Structural signal in SHAPE (NAI-N_3_) data for the 18S rRNA structure.

Supplemental Figure S10 Correlation between DMS and NAI-N_3_ probing of the mRNA secondary structure of the *psbA* translation initiation region.

Supplemental Figure S11 Footprint of a putative regulatory protein bound to the 5’ UTR of *psbA* and secondary structure of the *psbA* translation initiation region

Supplemental Figure S12 Reproducibility of translatomic and transcriptomic data.

Supplemental Figure S13 Changes in translation and mRNA levels of plastid-encoded genes.

Supplemental Figure S14 Correlations between changes in mRNA secondary structure and translation efficiency.

Supplemental Table 1 Number of mapped reads from the DMS-MaPseq analysis

Supplemental Table 2 Strength of binding of Shine-Dalgarno sequences to anti-Shine-Dalgarno sequences.

Supplemental Table 3 Fold change of mRNA levels and translation efficiency of nuclear-encoded genes encoding factors possibly regulating plastid translation.

Supplemental Table 4 List of used oligonucleotides used and their sequences.

## Conflict of Interest

No conflict of interest

## Author contributions

P.G., C.E., D.L., P.E.J, J.V., and L.B.S. conceived the study; P.G., C.E., P.K., and L.B.S performed experiments; S.M., S.K., P.E.J, and J.V. supervised experiments; P.G., C.E., J.V., and L.B.S. analyzed data; P.G., C.E., and L.B.S. wrote the article with contributions from all authors.

## Notes

### Competing Interest Statement

The authors have declared no competing interest.

