## Supplemental Data for "Light-dependent translation change of Arabidopsis *psbA* correlates with RNA structure alterations at the translation initiation region"

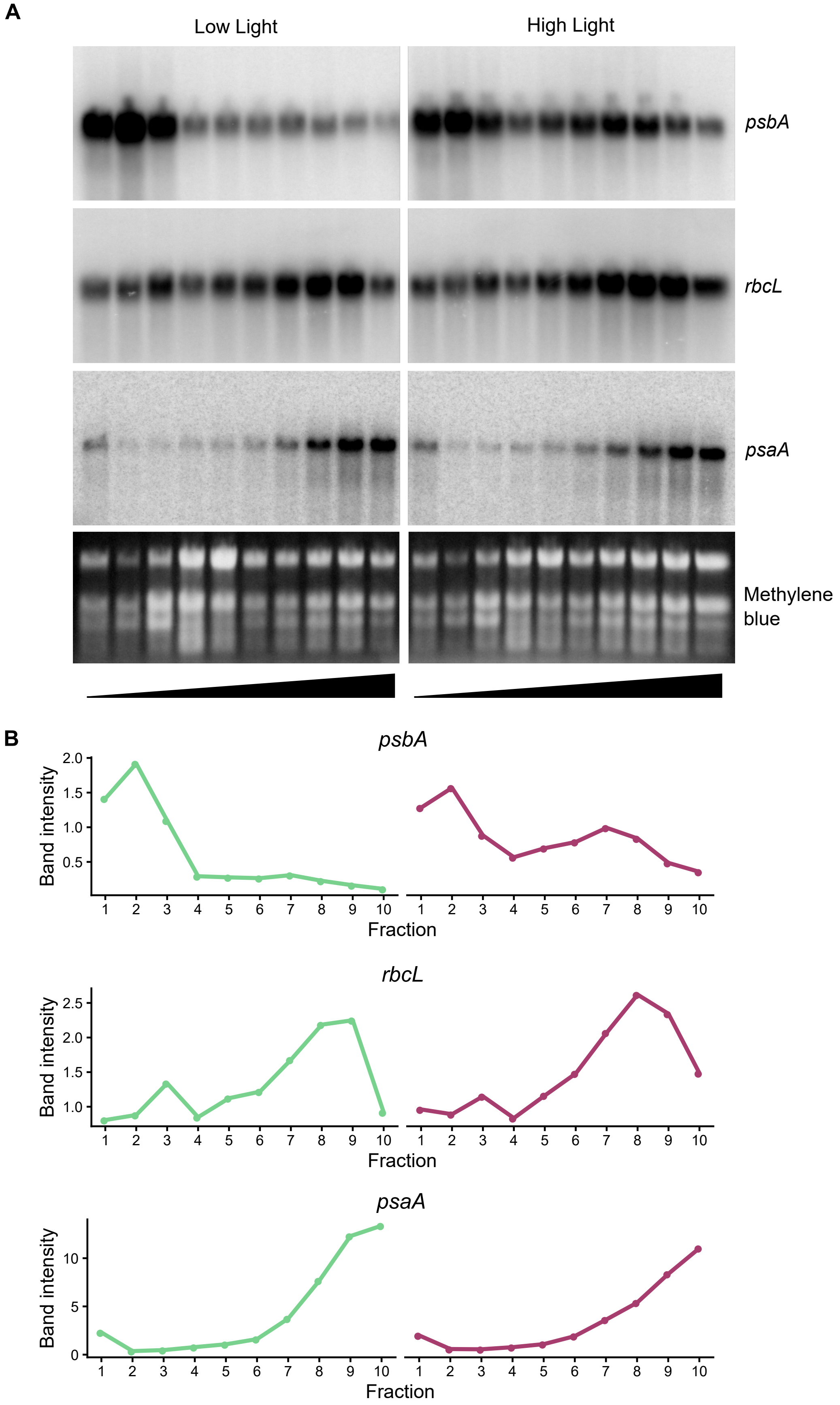


**Supplemental Figure S1 Polysome analysis of *psbA* and *rbcL* translation in young plants: A**, Analysis of *psbA* and *rbcL* translation in 17-18 days old plants after low light and high light treatment. *psaA* (encoding the PsaA subunit of photosystem I) is included as an example of a gene that is not upregulated in high light. Extracts from plants treated the same way as the plants analyzed by DMS-

MaPseq (Figure 1,3) were subjected to sucrose-gradient centrifugation, fractionated into ten fractions, and RNA was extracted from each. The upper panels show RNA gel blots probed for *psbA*, *rbcL*, and *psaA* mRNA, respectively. The lower panels show methylene blue-stained rRNAs. The gradients are schematically depicted as black triangles at the bottom: less dense fractions containing free mRNA are on the left, denser fractions containing polysome-associated mRNA actively engaged in translation are on the right. **B**, Quantification of the amount of *psbA*, *rbcL*, and *psaA* mRNA in each fraction normalized to the average amount of the five least dense fractions.

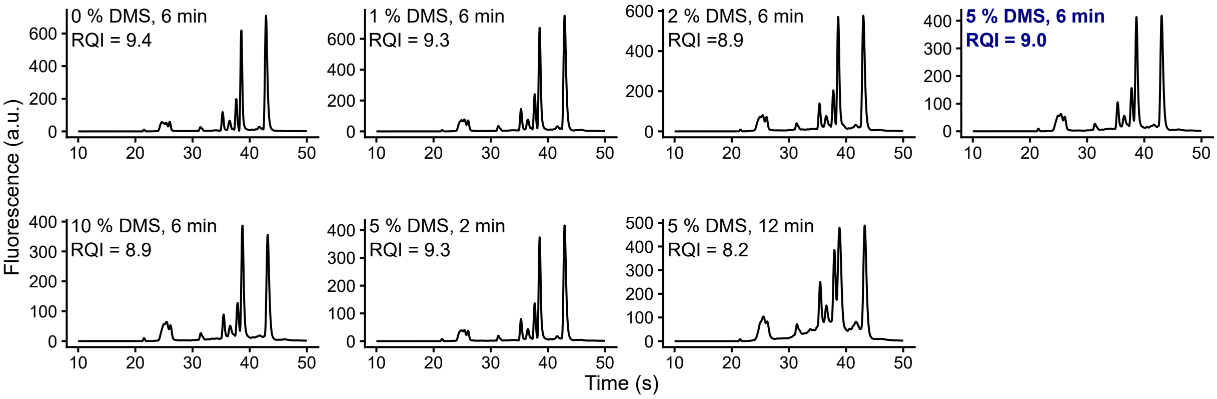


**Supplemental Figure S2 RNA quality after DMS treatment:** Quality assessment of RNA isolated after treatment with different DMS concentrations and different infiltration times. The results of capillary electrophoresis (Experion, Bio-Rad) are shown and the RNA Quality Index (RQI) is given (max = 10). Marked in bold and blue is the treatment that was used for RNA secondary structure probing (Figure 1 and 3).

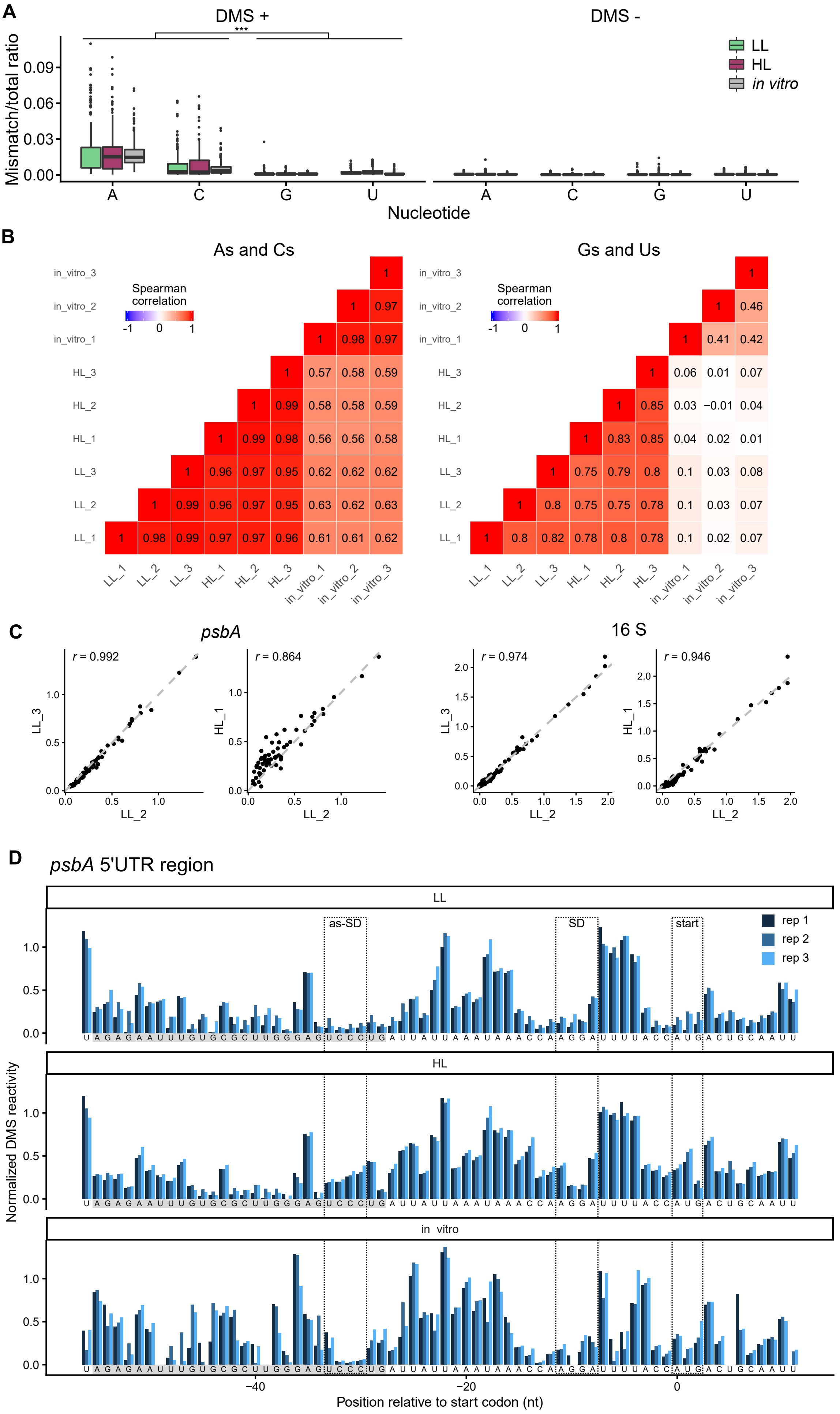


**Supplemental Figure S3 DMS-MaPseq quality control: A**, Mismatch rates shown for each nucleotide as determined by MaPseq of DMS-probed samples (DMS +) and the water control (DMS -). Probed nucleotides are detected by mismatches between the known mRNA sequence and the sequence obtained by DMS-MaPseq. Adenosines and cytidines are statistically more significantly probed than guanosines and uridines. (*P*-values calculated with the Tukey HSD test; *** = $P < 0.001$). For the coverage see Supplemental Table 1. **B**, Spearman correlations between replicates and samples. For low light (LL), high light (HL), and *in vitro*-folded RNA samples three biological replicates each were included. The left part shows the values for adenosines and cytidines, the right part for guanosines und uridines. Spearman's *r* values are shown both numerically and as color codes (see scale). **C**, Pairwise comparisons of the probing of adenosines and cytidines of *psbA* translation initiation region (left) and 16S rRNA (right) of representative sample pairs. Spearman's *r* value is given. **D**, Normalized DMS reactivities for *psbA* in low light (LL), high light (HL), and *in vitro*-folded RNA samples with three biological replicates each. The positions of a sequence (as-SD) that can bind the Shine-Dalgarno sequence, the Shine-Dalgarno sequence (SD), and the start codon (start) are marked with dashed boxes (see Figure 1). The footprint of an RNA binding protein (Supplemental Figure S11A-D) is indicated in the sequence with a gray background.

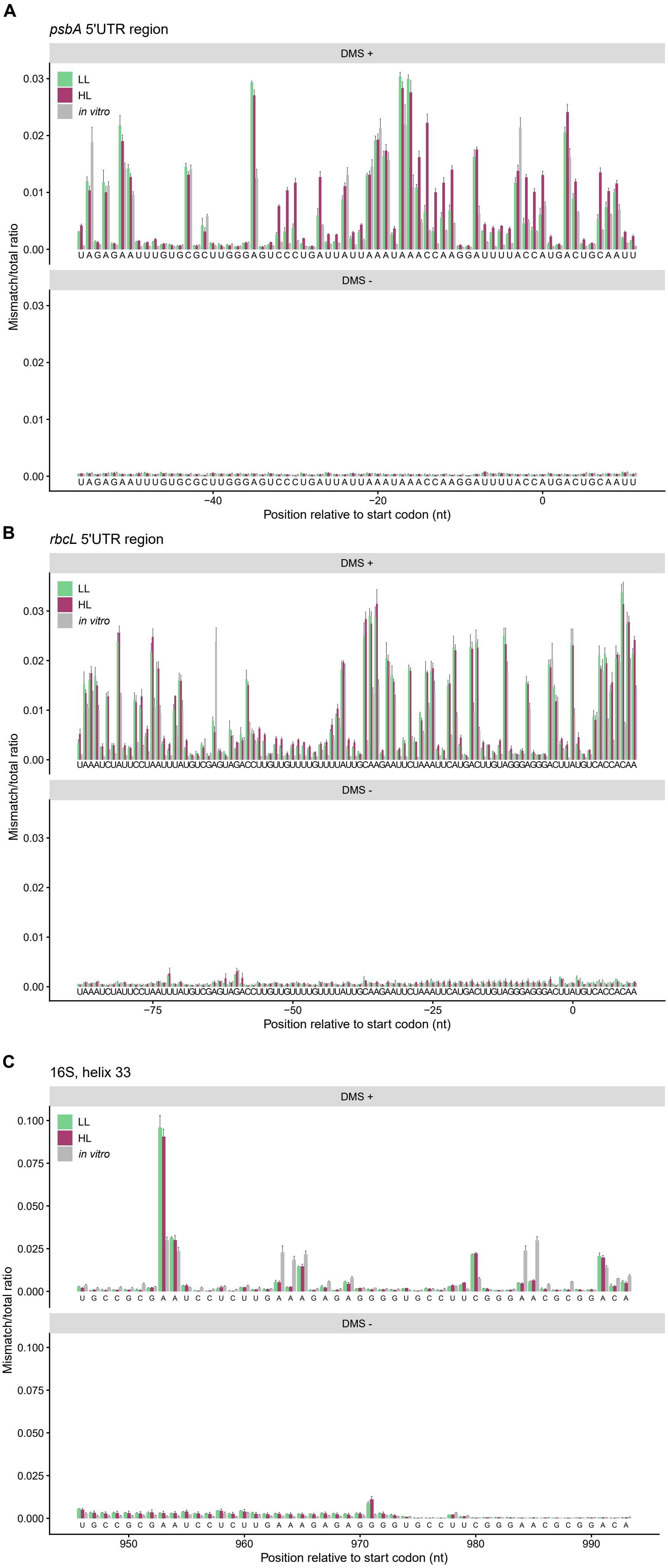


**Supplemental Figure S4 DMS-MaPseq compared to water control**: **A**, DMS-based mapping of secondary structure of the *psbA* translation initiation region of 17-18-days-old plants (DMS +) compared to water control (DMS -) as determined by MaPseq. Control experiment for Figure 1A. The data for the low light (LL) control are shown in light green, for the high light (HL) sample in dark red, and for the mRNA that was allowed to fold *in vitro* in gray. **B**, Water control for DMS-MaPseq of the *rbcL* translation initiation region (Figure 3A). **C**, Water control for DMS-MaPseq of helix 33 of the 16S rRNA. Average values can be found in Supplemental Figure S3A.

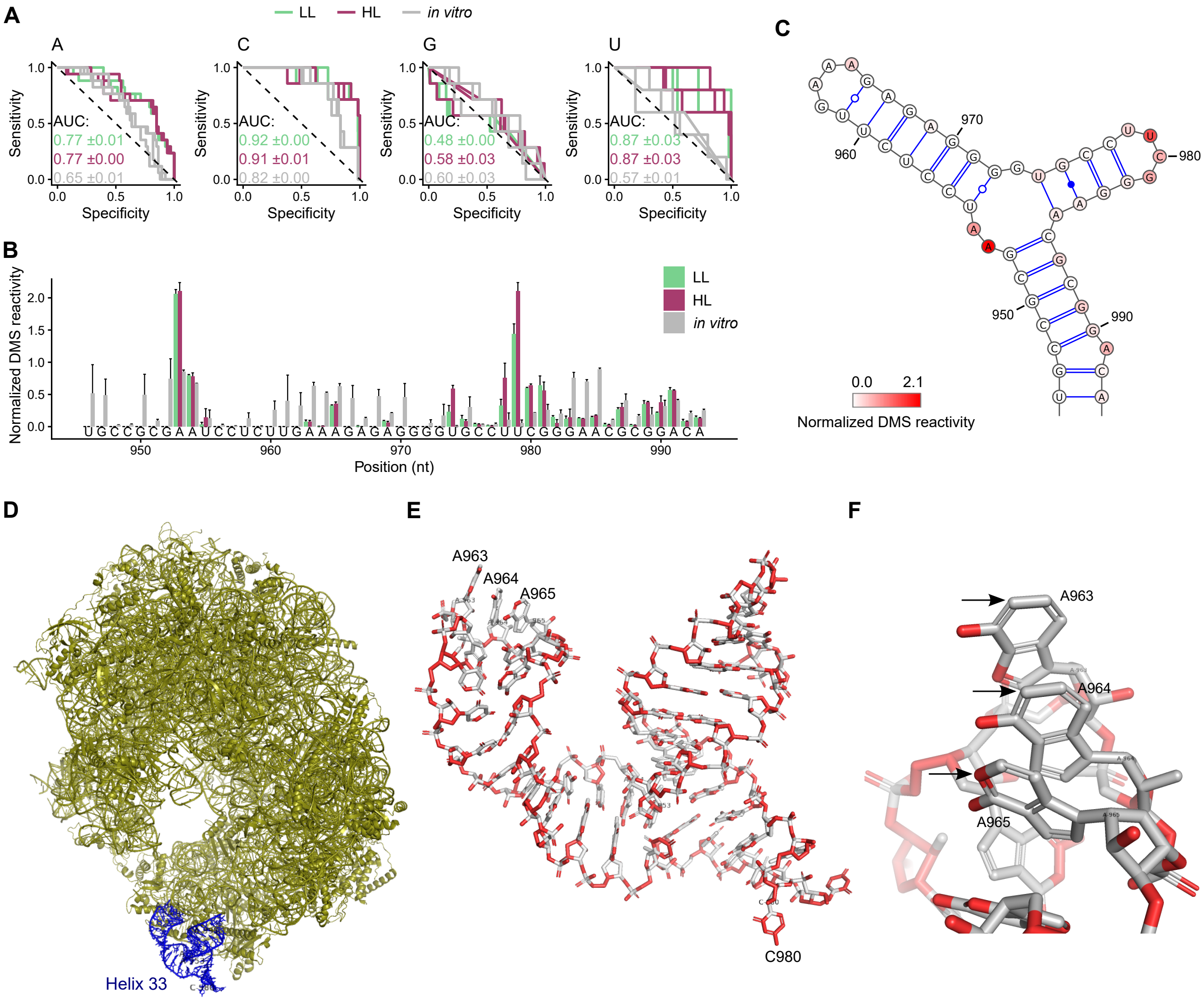


**Supplemental Figure S5 DMS data reproduces known elements of the secondary structure of 16S rRNA**: The secondary structure of helix 33 of the plastid 16S rRNA was analyzed by probing it with DMS. **A**, Receiver Operating Characteristics (ROC) curve generated using the Arabidopsis 16S rRNA secondary structure (Ahmed et al., 2017) as predictor and the normalized DMS reactivities as response. The ROC curves were calculated separately for each nucleotide (compare to Supplemental Figure S3A) and for each of the three biological replicates of each analyzed condition. Light green is the low light (LL) control, dark red the high light (HL) treatment, and gray RNA that was allowed to fold *in vitro*. From the generated ROC curve, the average area under curve (AUC) was calculated for each analyzed condition. As expected (Mustoe et al., 2019; Gawroński et al., 2020), the structure signal is better for adenosines and cytidines than for guanosines and uridines. In addition, the *in vivo* DMS reactivities (LL, HL) reproduce the structure better than the ones at the *in vitro*-folded, protein-free 16S rRNA. **B**, Helix 33 was analyzed in more detail. High normalized DMS reactivity indicates high levels of bound probe, i.e. single-stranded regions. The *in-vivo* rRNA structure was not altered by the high light treatment, whereas there are multiple differences between the *in-vivo*, protein-bound 16S rRNA (LL and HL samples) and the *in-vitro*, protein-free 16S rRNA. **C**, The low-light data shown in **B** are here depicted as a simplified RNA structure based on the previously published structure of the plastid ribosome (Ahmed et al., 2017). DMS values for LL and HL samples fit the published ribosome structure well, with the exception of those for A963 and A964. **D**, Structure of the 16S rRNA (Ahmed et al., 2017), with helix 33 marked in blue. **E**, Magnification of helix 33 with adenosines A963-A965 and cytidine C980 indicated. **F**, The accessibility of each atom in the loop was predicted as described in the Methods section; accessible atoms are marked in red. The atoms that should be susceptible to methylation by DMS are marked with arrows. The poor probing of A963 and A964 by DMS is not caused by their involvement in base pairing but by their (relative) inaccessibility to DMS.

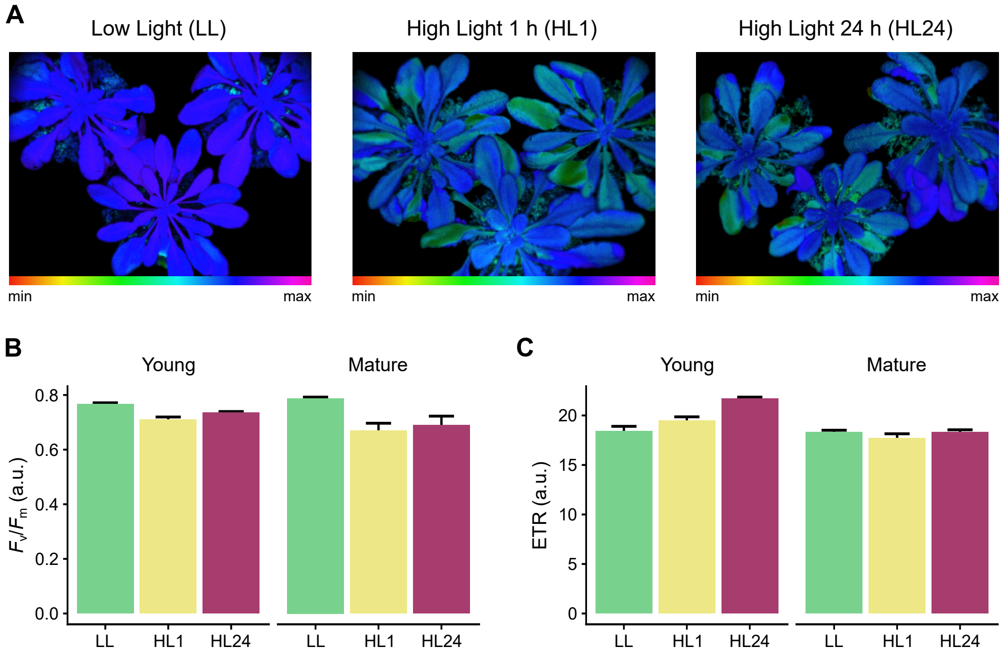


**Supplemental Figure S6 Determination of photosynthetic parameters**: **A**, False colored images of 7-week-old plants exposed to different light treatments showing the maximum quantum yields of photosystem II ($F_v/F_m$). Plants grown under control low light conditions (LL; 140-160 µE $m^{-2}$ $s^{-1}$), after 1 h of high light (HL1; 1200 µE $m^{-2}$ $s^{-1}$), and after 24 h of high light (HL24; 4 h, 1200 µE $m^{-2}$ $s^{-1}$; 16 h dark; 4 h, 1200 µE $m^{-2}$ $s^{-1}$) are compared. **B**, Average $F_v/F_m$ values for young (not fully expanded) and mature (fully expanded) leaves of three plants for each condition shown in **A**. **C**, Average electron transport rate (ETR). Values are given in arbitrary units (a.u.). Error bars indicate mean standard errors.

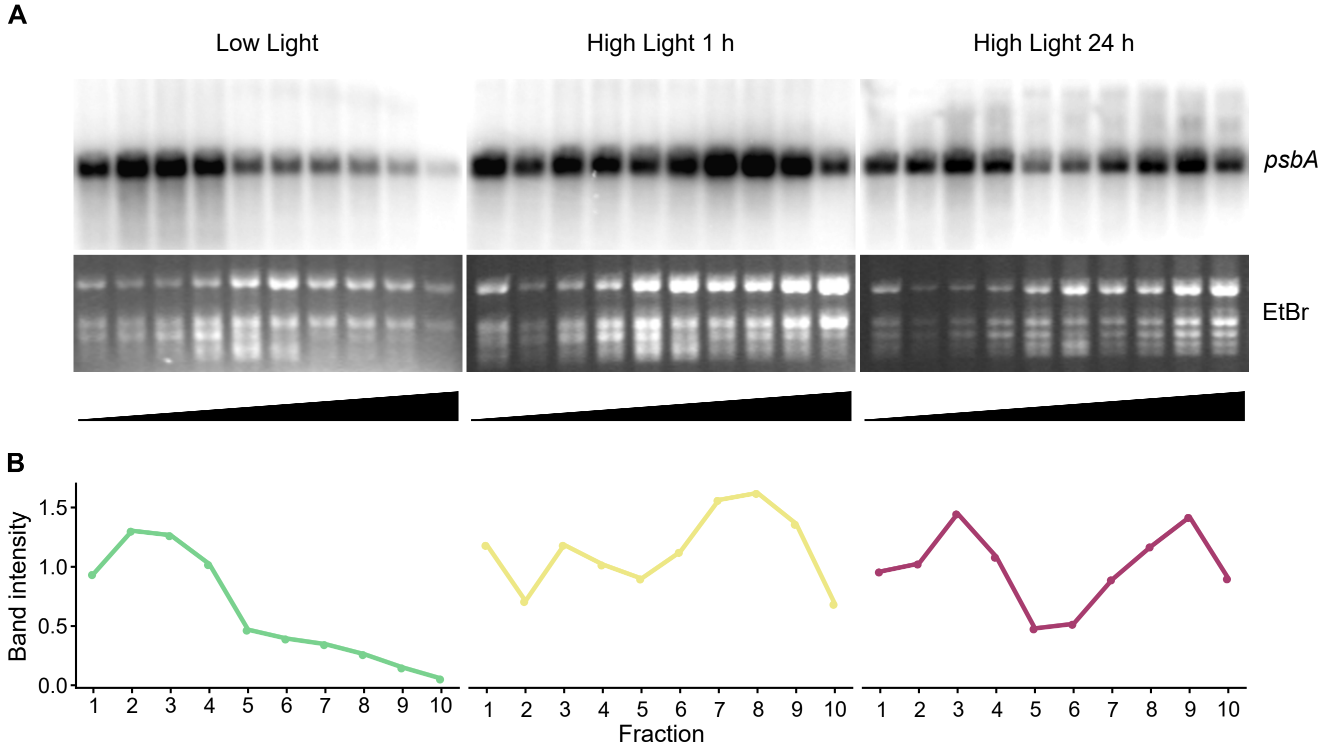


**Supplemental Figure S7 Polysome analysis of *psbA* translation in mature plants**: Comparison of low light control, 1 h high light and 24 h high light treatment (4 h high light, 16 h dark, 4 h high light). The low light control and the 24 h treatment were identical to the conditions used for the SHAPE experiment (Figure 1,3) and ribosome profiling (Figure 4). Plant extracts were subjected to sucrose-gradient centrifugation, fractionated into ten fractions, and RNA was extracted from each. The upper panels show RNA gel blots probed for *psbA* mRNA. The lower panels show ethidium bromide (EtBr)-stained rRNAs. The gradients are schematically depicted as black triangles at the bottom: less dense fractions containing free *psbA* mRNA are on the left, denser fractions containing polysome-associated *psbA* mRNA actively engaged in translation are on the right. **B**, Quantification of the amount of *psbA* mRNA normalized to the average amount of the five least dense fractions.

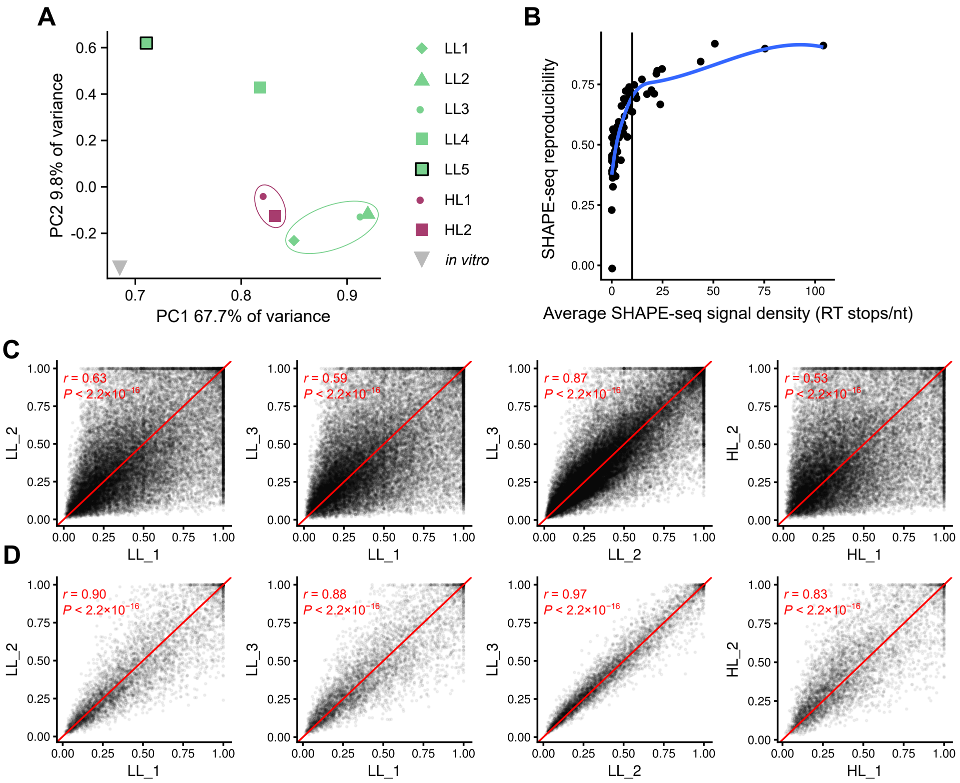


**Supplemental Figure S8 Reproducibility of the mRNA secondary structure probing (NAI-$N_3$) data**: **A**, Principal component analysis (PCA). The low light (LL) samples are shown in light green, the high light (HL) samples in dark red, and the sample that was folded *in vitro* is in gray. LL1 and LL2 are technical replicates separated after DNase treatment; the other samples are true biological replicates. Samples LL3, LL5, HL1, HL2, and the *in vitro* sample were depleted of rRNA prior to probing; the other samples are total RNA samples. Samples LL4 and LL5 were excluded from further analysis because of lack of similarity both to the other samples and to each other. The samples chosen for further analysis are encircled. **B**, Reproducibility is dependent on the average signal intensity (number of reverse transcription (RT) stops per nucleotide). The vertical line indicates the chosen cutoff of ≥10 reverse transcription stops/nt on average for each gene. Each point represents the value for one plastid-encoded gene. **C**, Pairwise comparisons of the three low light (LL1-LL3) and two high light samples (HL1, HL2). All data is represented, i.e. no cutoff is used. Each point represents the data for one nucleotide of a plastid mRNA. Spearman's *r* and *P* values are given. **D**, Pairwise comparisons of the same samples as in C but with a cutoff of ≥10 reverse transcription stops/nt on average for each gene.

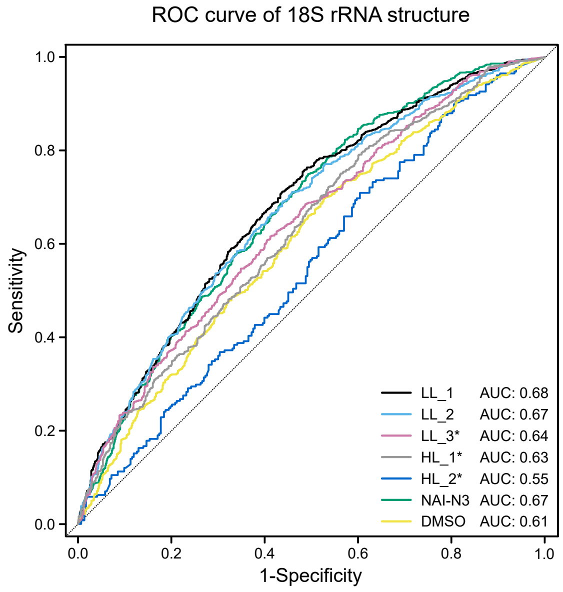


**Supplemental Figure S9 Structural signal in SHAPE (NAI-$N_3$) data for the 18S rRNA structure**: Receiver Operating Characteristics (ROC) curve generated using the Arabidopsis 18S rRNA secondary RNA structure from the CRW data as predictor (Cannone et al., 2002) and the swinsor-normalized termination counts as response. From the generated ROC curve, the area under curve (AUC) was calculated and indicated in the legend for each tested sample. Three low light samples (LL1-LL3) and two high light samples (HL1, HL2) are shown. A control without the biotin-based selection step (NAI-$N_3$) and a control treated only with DMSO are also included. The samples LL3, HL1, and HL2 (indicated with asterisks) were depleted of rRNA and had significantly lower coverage of the 18S rRNA, resulting in lower RNA structure probing signals.

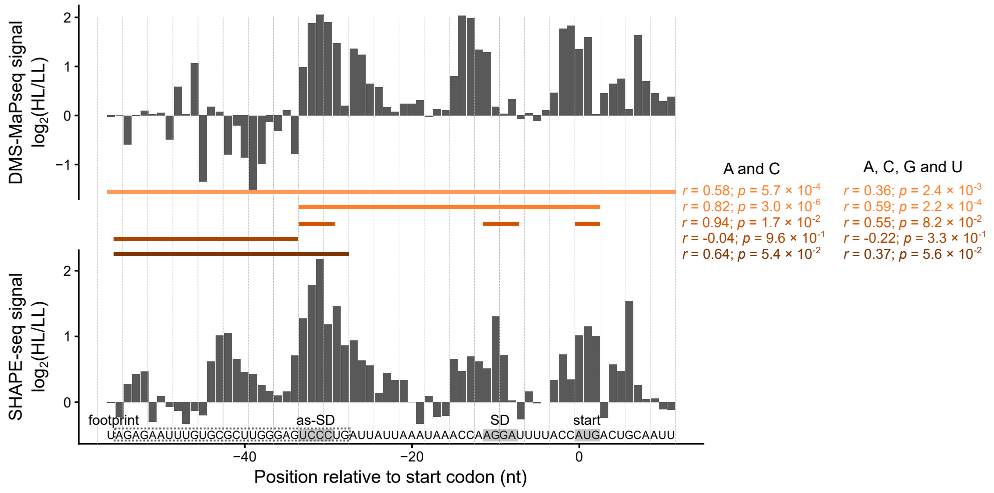


**Supplemental Figure S10 Correlation between DMS and NAI-N$_3$ probing of the mRNA secondary structure of the *psbA* translation initiation region:** Correlation analysis of SHAPE and DMS probing (Figure 1A,C). Shown are the ratios of the high light (HL) to the low light (LL) signals, expressed as $\log_2$ values. Spearman's $r$ with the corresponding $p$ values for the correlation of the SHAPE and DMS data for different parts of the *psbA* translation initiation region are given; the corresponding regions are indicated above the sequence in the same color. The correlation was analyzed separately for the more reliable probing of adenosines and cytidines (Supplemental Figures S3B, S5B) and for all four nucleotides. The structural data is well correlated at the *cis*-elements important for translation initiation but less well at the footprint (Figure S11A) excluding the sequence that can bind the SD (as-SD). This could be caused by differences between NAI-N$_3$ and DMS in the sensitivity to protein binding. DMS probing is sensitive to protein binding (Kwok et al., 2013; Talkish et al., 2014), whereas SHAPE reagents provide information about structural changes caused by protein binding, but bound proteins are not always detected as nucleotides with low reactivity (Spitale et al., 2013, 2015; Kenyon et al., 2015).

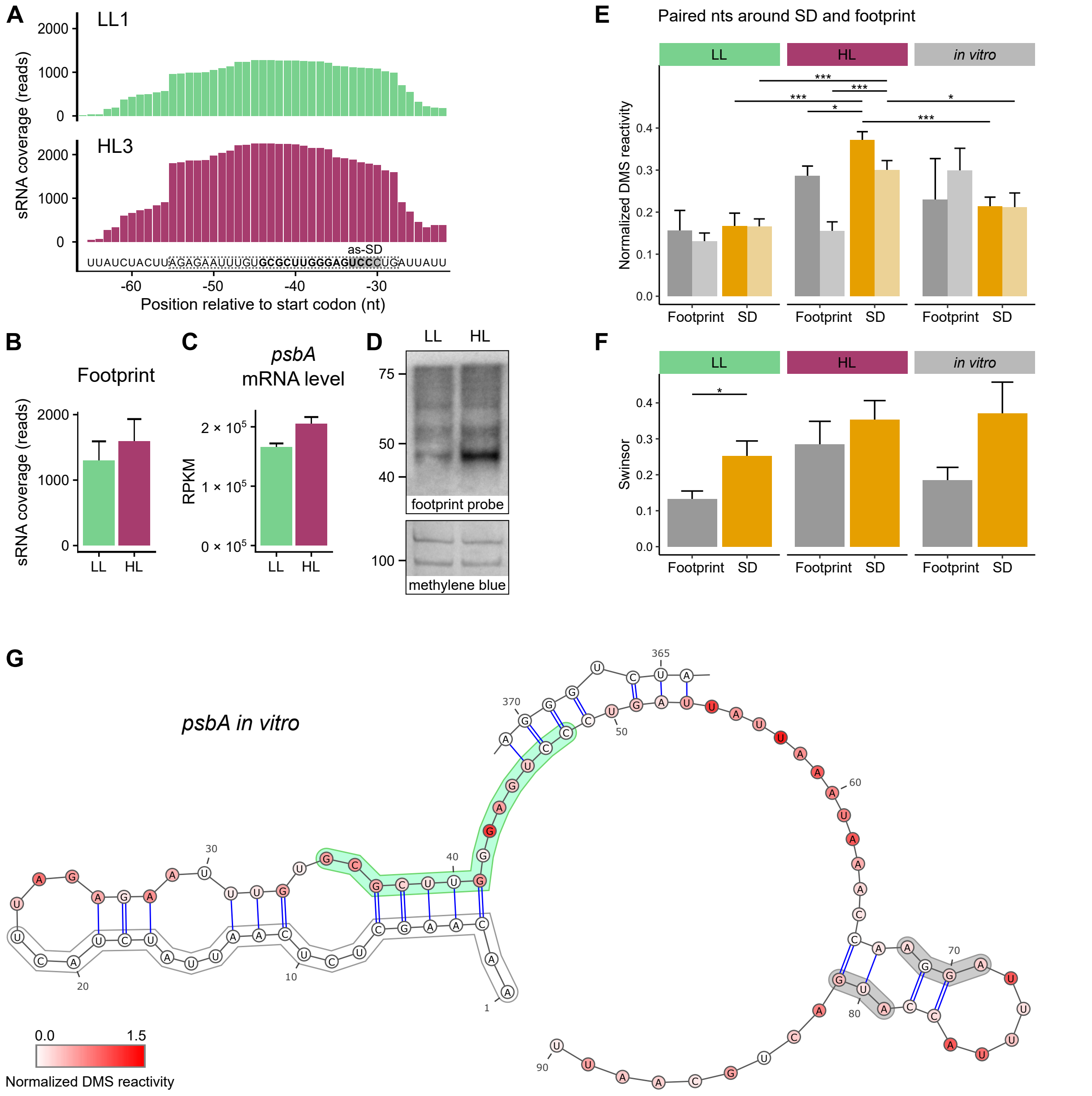


**Supplemental Figure S11 Footprint of a putative regulatory protein bound to the 5' UTR of *psbA* and secondary structure of the *psbA* translation initiation region**: **A**, The number of reads detected in the region of the footprint are shown for one example each of a low light control (LL1) and a high light treatment (HL3). MNase-digested RNA samples from 7-week-old plants were analyzed. The dashed box shows the position of the footprint found in this analysis, the bold nucleotides indicate the footprint of HCF173 (plus possibly other proteins) found earlier (McDermott et al., 2019), the sequence that can bind the Shine-Dalgarno sequence (as-SD) is marked in gray. **B**, Average coverage of the footprint in three biological replicates each for the low light control (light green) and high light samples (dark red). **C**, *psbA* mRNA levels in low light (light green) and after exposure to high light (dark red). Reads per kilobase of transcript per million mapped reads (RPKM) values are shown. **D**, Footprint detected in low light and high light samples by RNA gel blot analysis using a probe specific for the footprint. The same plant material as in **A** was used. The isolated RNA was not treated with any RNase.

Methylene blue-stained rRNAs are shown in the bottom panel as loading controls. Note that the footprint differs in size. It is shortest after RNase I treatment (McDermott et al., 2019) and longest when analyzed without any additional RNase treatment (**D**). **E**, Average normalized DMS reactivities of the nucleotides predicted to form base pairs in low light (Figure 2A) shown separately for the two halves of the stem loop: the footprint side (between nucleotides 35-48) and the SD side (between nucleotides 69-86) (compare Figure 2C). Values for low light (LL), high light (HL), and *in vitro*-folded RNA are shown. The darker colored columns represent the more reliable reactivities at adenosines/cytidines, the columns in lighter colors are the reactivities at all four nucleotides. Note also the sequence bias, especially the enrichment cytidines near the 3' end of the footprint (Figure 2A). Asterisks here (and in **F**) indicate statistically significant changes (*P*-values calculated with the Wilcoxon rank sum test; * = $P < 0.05$, ** = $P < 0.01$, and *** = $P < 0.001$), error bars indicate mean standard error. **F**, Average normalized SHAPE reactivities (swinsor) of the nucleotides predicted to form base pairs in low light (Figure 2A) shown separately for the two halves of the stem loop: the footprint side and the SD side. **G**, Predicted mRNA secondary structure of the *in vitro*-folded RNA using normalized DMS reactivities (Figure 1A) as constrains (control for Figure 2A,B). The prediction includes an interaction with a part of the coding region (position 364-371).

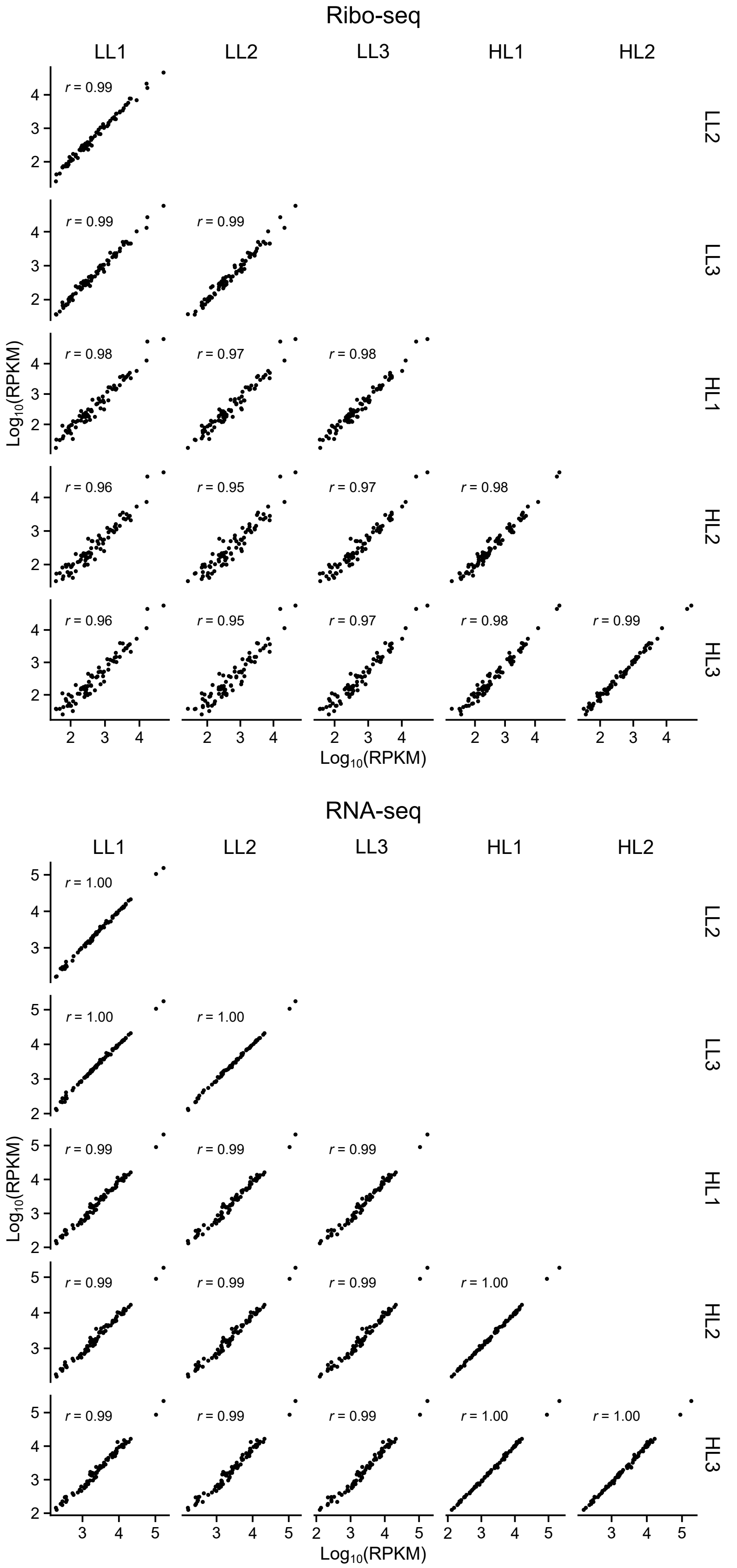


**Supplemental Figure S12 Reproducibility of translatomic and transcriptomic data**: Pairwise comparison of the three replicates for both low light (LL) and high light (HL) (24 h). The upper part contains the translatomic (ribosome profiling data, Ribo-seq) data, the lower one the transcriptomic (RNA-seq) data. Each set of points represents the $Log_{10}$ Reads per kilobase of transcript per million mapped reads (RPKM) values for one plastid-encoded coding region. Spearman's *r* value is given. The response to the high light treatment occurs primarily at the level of translation (see also Figure 4 and Supplemental Figure S13).

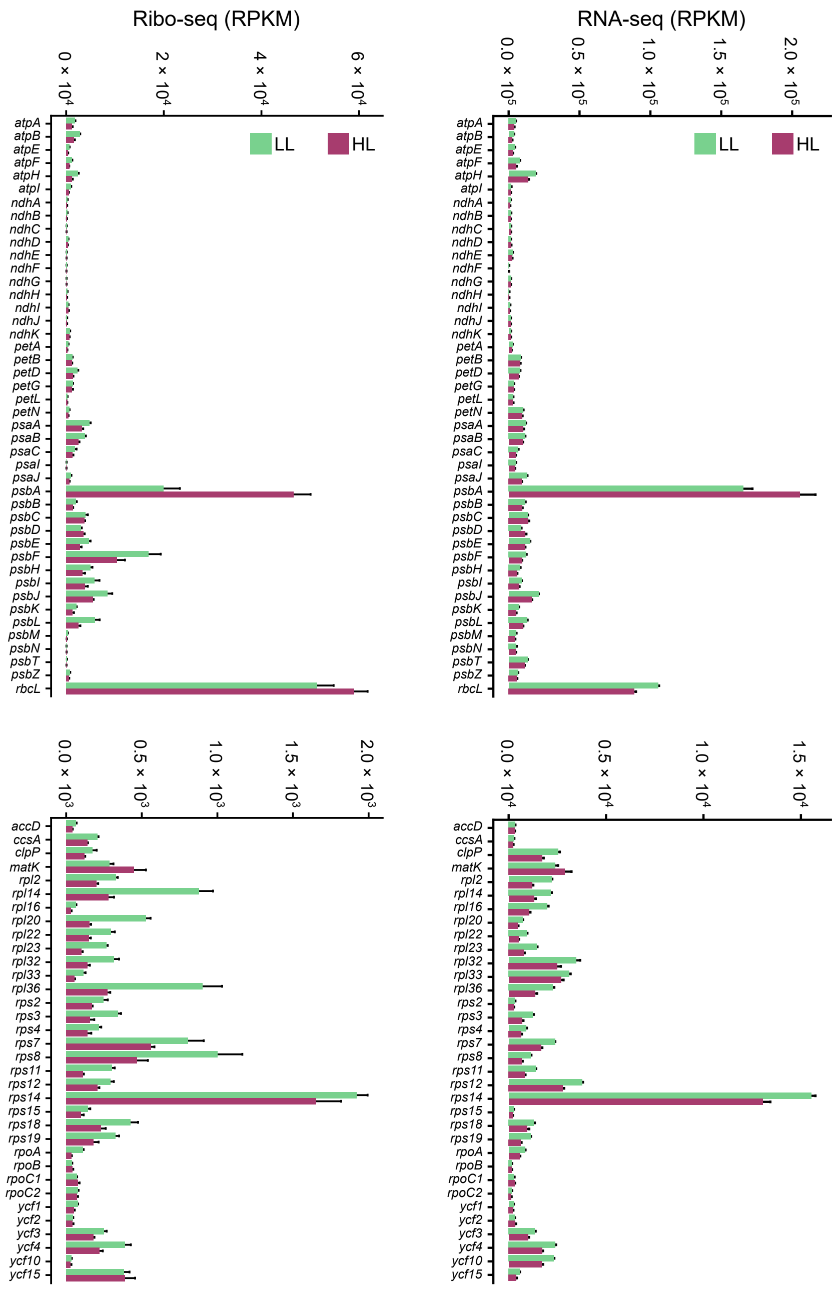


**Supplemental Figure S13 Changes in translation and mRNA levels of plastid-encoded genes**: Reads per kilobase of transcript per million mapped reads (RPKM) values are shown for each gene. The left panels depict the ribosome profiling data (Ribo-seq), the right panels show mRNA levels (RNA-seq). The values from the low light (LL) control are shown in light green, those from the 24h high light (HL) treatment in dark red. The data in the upper panels are derived from all genes coding for subunits of the photosynthetic complexes, the lower panels show the results for all other genes. Compare to the translation efficiency (Ribo-seq/RNA-seq) in Figure 4 (see also Figure S12 for data reproducibility).

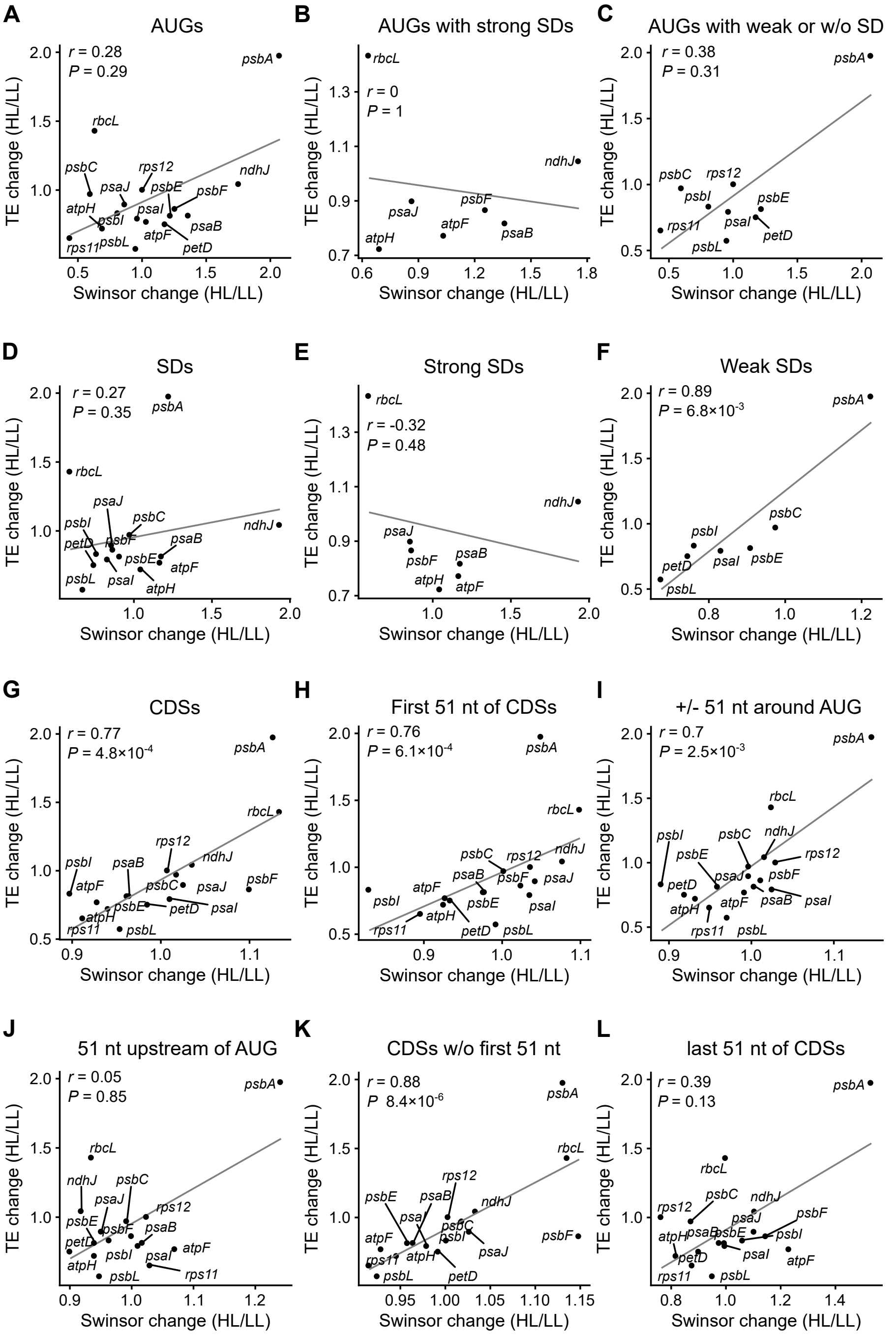


**Supplemental Figure S14 Correlations between changes in mRNA secondary structure and translation efficiency**: The changes in mRNA secondary structure are calculated from the swinsor-normalized termination count values derived from NAI-$N_3$ probing by dividing the values for the plants exposed to high light (HL) by those obtained from the low light (LL) controls. An increase in the swinsor value reflects a decrease in base pairing, i.e. less RNA secondary structure. Average changes for the indicated region of each gene are given. The change in translation efficiency is calculated by dividing the normalized read counts for the ribosomal footprints by the normalized read counts for the transcripts for each coding region, and then dividing the obtained values from the high light treatment by those from the low light control. Only genes with sufficient coverage of the mRNA secondary structure (in average at least 10 reverse transcription stops/nucleotide) are included. Spearman's *r* and *P* values are given. The correlation of the change of translation efficiency and the change of RNA secondary structure at the following regions is shown: **A**, start codons (AUGs); **B**, start codons of genes with strong Shine-Dalgarno sequences (SD) (hybridization to the anti-SD of the 16S rRNA < -9 kcal $mol^{-1}$); **C**, start codons of genes with weak or without SD (> -6 kcal $mol^{-1}$); **D**, SDs; **E**, SDs of genes with strong SD (< -9 kcal $mol^{-1}$); **F**, SDs of genes with weak SD (> -6 kcal $mol^{-1}$); **G**, complete coding regions (CDSs); **H**, the first 51 nt of the coding regions including the start codons; **I**, the 51 nt around start codons; **J**, the 51 nt of the 5' UTRs upstream of the start codons; **K**, coding regions without the first 51 nt, and **L**, the last 51 nt of the coding regions. We assume that, as described for *E. coli* (Mustoe et al., 2018), the correlation in the case of the CDS (G, K) is a consequence of differences in translation initiation, and that the structural changes of the CDS are caused by translating ribosomes reducing the mRNA secondary structure in the coding region.

**Supplemental Table 1 Number of mapped reads from the DMS-MaPseq analysis**. Shown are the values for the three biological replicates for in vitro folded RNA, low light (LL), and high light (HL); both for the samples including DMS (DMS+) and the controls without DMS (DMS-).

| **sample name** | ***psbA*** | **16S** | ***rbcL*** |
|---|---|---|---|
| *in vitro* 1 DMS+ | 59378 | 30168 | 11662 |
| *in vitro* 2 DMS+ | 57866 | 25334 | 9423 |
| *in vitro* 3 DMS+ | 68688 | 31449 | 14247 |
| *in vitro* 1 DMS- | 64056 | 43671 | 11989 |
| *in vitro* 2 DMS- | 60806 | 41152 | 13989 |
| *in vitro* 3 DMS- | 47834 | 31732 | 12155 |
| LL 1 DMS+ | 51107 | 37692 | 16024 |
| LL 2 DMS+ | 87031 | 39773 | 15312 |
| LL 3 DMS+ | 57313 | 34316 | 15947 |
| LL 1 DMS- | 62580 | 31825 | 18332 |
| LL 2 DMS- | 52764 | 29991 | 16549 |
| LL 3 DMS- | 45224 | 27490 | 13855 |
| HL 1 DMS+ | 73896 | 27997 | 14164 |
| HL 2 DMS+ | 61158 | 37134 | 13469 |
| HL 3 DMS+ | 67462 | 39944 | 17238 |
| HL 1 DMS- | 82393 | 31700 | 21708 |
| HL 2 DMS- | 56735 | 35606 | 15266 |
| HL 3 DMS- | 67588 | 39983 | 15526 |

**Supplemental Table 2 Strength of binding of Shine-Dalgarno sequences to the anti-Shine-Dalgarno sequence:** The strength of the hybridization of the nucleotides -22 to -2 of each 5' UTR to the anti-Shine-Dalgarno sequence of the 16S rRNA was calculated *in silico* using Free2bind (Starmer et al., 2006) for all 16 genes whose RNA secondary structure could be analyzed. As comparison, the *psbA* and *rbcL* genes of *Nicotiana tabacum* and *Zea mays* are included. For each species, a typical growth temperature was used for the calculations (20°C for Arabidopsis, 22°C for *N. tabacum*, and 31°C for *Z. mays*).

| Species | Gene | SD-aSD binding [kcal/mol] | |
|---|---|---|---|
| *Arabidopsis thaliana* | *rbcL* | -12,98 | strong SDs |
| | *psaJ* | -12,94 | |
| | *psaB* | -12,84 | |
| | *psbF* | -12,84 | |
| | *atpF* | -9,74 | |
| | *atpH* | -9,28 | |
| | *ndhJ* | -9,24 | |
| | *psaI* | -5,80 | Weak SDs |
| | *psbA* | -5,50 | |
| | *psbE* | -5,20 | |
| | *psbI* | -5,20 | |
| | *petD* | -4,84 | |
| | *psbL* | -1,40 | |
| | *psbC* | -0,84 | |
| | *rps11* | 1,14 | No SD |
| | *rps12* | 1,14 | |
| *Zea mays* | *rbcL* | -11,23 | |
| | *psbA* | 1,51 | |
| *Nicotiana tabacum* | *rbcL* | -12,66 | |
| | *psbA* | 1,21 | |

**Supplemental Table 3** Fold change of mRNA levels and translation efficiency of nuclear-encoded genes encoding factors possibly regulating plastid translation: $Log_2$ values of the fold change in high light samples relative to the low light controls and the corresponding p values are given.

| Gene | Name | mRNA levels $log_2$ fold change | p value | Translation efficiency $log_2$ fold change | p value |
|---|---|---|---|---|---|
| AT4G16390 | ATP4, SVR7 | -0,27 | 1,00E+00 | -0,50 | 3,56E-01 |
| AT1G77510 | AtPDI6 | 0,16 | 1,00E+00 | -0,16 | 7,51E-01 |
| AT5G42310 | CRP1 | 0,13 | 1,00E+00 | -0,08 | 6,97E-01 |
| AT3G46790 | CRR2 | 0,60 | 1,00E+00 | 0,21 | 5,29E-01 |
| AT3G17040 | HCF107 | 0,04 | 1,00E+00 | -0,15 | 6,10E-01 |
| AT5G08720 | HCF145 | 0,70 | 1,00E+00 | -0,17 | 6,18E-01 |
| AT3G09650 | HCF152 | 1,04 | 1,00E+00 | 0,16 | 5,01E-01 |
| AT1G16720 | HCF173 | 0,03 | 1,00E+00 | 0,08 | 8,38E-01 |
| AT4G35250 | HCF244 | 0,50 | 1,00E+00 | -0,35 | 1,57E-01 |
| AT3G46610 | LPE1 | 0,19 | 1,00E+00 | -0,08 | 7,77E-01 |
| AT4G34830 | MRL1 | 0,45 | 1,00E+00 | 0,16 | 5,29E-01 |
| AT5G02120 | OHP1 | 0,28 | 1,00E+00 | -0,22 | 3,83E-01 |
| AT1G34000 | OHP2 | 0,27 | 1,00E+00 | -0,60 | 1,17E-01 |
| AT1G71720 | PBR1, PDE338, RLSB | -0,11 | 1,00E+00 | 0,20 | 6,18E-01 |
| AT4G31850 | PGR3 | 0,14 | 1,00E+00 | 0,15 | 5,03E-01 |
| AT2G18940 | PPR10 | 0,15 | 1,00E+00 | -0,11 | 7,67E-01 |
| AT5G46580 | PPR53 | 0,42 | 1,00E+00 | -0,14 | 7,66E-01 |

**Supplemental Table 4 List of used oligonucleotides and their sequences.** Blue are indexes.

| Name | Sequence (5' --> 3') | Used for |
|---|---|---|
| psbAfor | AAGCGAAAGCCTATGGGGTC | probe for polysomes |
| psbArev | AATGTTGTGCTCAGCCTGGA | probe for polysomes |
| rbcLfor | GGACAACTGTGTGGACCGAT | probe for polysomes |
| rbcLrev | ATACCGCGGCTTCGATCTTT | probe for polysomes |
| psbA_footprint_probe | AGGGACTCCCAAGCGCACAAATTCTCT | probe for small RNA blot |
| DNA linker | CTGTAGGCACCATCAAT | determination of 3'-ends of plastid transcripts |
| Linker primer | 5Phos/ATCTCGTATGCCGTCTTCTGCTTG/ iSp18/CACTCA/ iSp18/TCCGACGATCATTGATGGTGCCTACAG | determination of 3'-ends of plastid transcripts |
| RRN16S_1048_MaP_R1 | CGGGACTTAACCCAACACC | RNA structure probing with DMS |
| psbA_MaP_RP | TGTAGATGGAGCCTCAACAGCA | RNA structure probing with DMS |
| rbcL_MaP_RP | AGATTGAGCCGAGTGCAATTAAACT | RNA structure probing with DMS |
| 16S_540_MaP_IL_F | ACACGTTCAGAGTTCTACAGTCCGACGATCTTTAAGTCCGCCGTCAAATC | RNA structure probing with DMS |
| 16S_1048_MaP_IL_R2 | GAGTTCCTTGGCACCCGAGAATTCCA CGGGACTTAACCCAACACC | RNA structure probing with DMS |
| psbA_1_MaP_ILMN_F | ACACGTTCAGAGTTCTACAGTCCGACGATCAACAAGCTCTCAATTATCTACT | RNA structure probing with DMS |
| psbA_174_MaP_ILMN_R2 | GAGTTCCTTGGCACCCGAGAATTCCA CCATCCAATGTAAAGACGGT | RNA structure probing with DMS |
| rbcL_1_MaP_IL_F | ACACGTTCAGAGTTCTACAGTCCGACGATCGTATTTGGCGAATCAAATATCATGGTCT | RNA structure probing with DMS |
| rbcL_578_MaP_IL_R | GAGTTCCTTGGCACCCGAGAATTCCACGTAGAGCAGCCAGGGCTTT | RNA structure probing with DMS |
| ILMN_PCR1 | AATGATACGGCGACCACCGAGATCTACACGTTCAGAGTTCTACAGTCCGACGATC | RNA structure probing with DMS |
| ILMN_PCR2i1 | CAAGCAGAAGACGGCATACGAGAT **CGTGAT**GTGACTGGAGTTCCTTGGCACCCGAGAATTCCA | RNA structure probing with DMS, in_vitro_1 DMS+ |
| ILMN_PCR2i3 | CAAGCAGAAGACGGCATACGAGAT **GCCTAA**GTGACTGGAGTTCCTTGGCACCCGAGAATTCCA | RNA structure probing with DMS, in_vitro_2 DMS+ |
| ILMN_PCR2i8 | CAAGCAGAAGACGGCATACGAGAT **TCAAGT**GTGACTGGAGTTCCTTGGCACCCGAGAATTCCA | RNA structure probing with DMS, in_vitro_3 DMS+ |
| ILMN_PCR2i9 | CAAGCAGAAGACGGCATACGAGAT **CTGATC**GTGACTGGAGTTCCTTGGCACCCGAGAATTCCA | RNA structure probing with DMS, in_vitro_1 DMS- |
| ILMN_PCR2i10 | CAAGCAGAAGACGGCATACGAGAT **AAGCTA**GTGACTGGAGTTCCTTGGCACCCGAGAATTCCA | RNA structure probing with DMS, in_vitro_2 DMS- |
| ILMN_PCR2i11 | CAAGCAGAAGACGGCATACGAGAT **GTAGCC**GTGACTGGAGTTCCTTGGCACCCGAGAATTCCA | RNA structure probing with DMS, in_vitro_3 DMS- |
| ILMN_PCR2i13 | CAAGCAGAAGACGGCATACGAGAT **TTGACT**GTGACTGGAGTTCCTTGGCACCCGAGAATTCCA | RNA structure probing with DMS, LL_1 DMS+ |
| ILMN_PCR2i14 | CAAGCAGAAGACGGCATACGAGAT **GGAACT**GTGACTGGAGTTCCTTGGCACCCGAGAATTCCA | RNA structure probing with DMS, LL_2 DMS+ |
| ILMN_PCR2i15 | CAAGCAGAAGACGGCATACGAGAT **TGACAT**GTGACTGGAGTTCCTTGGCACCCGAGAATTCCA | RNA structure probing with DMS, LL_3 DMS+ |
| ILMN_PCR2i16 | CAAGCAGAAGACGGCATACGAGAT **GGACGG**GTGACTGGAGTTCCTTGGCACCCGAGAATTCCA | RNA structure probing with DMS, LL_1 DMS- |
| ILMN_PCR2i17 | CAAGCAGAAGACGGCATACGAGAT **CTCTAC**GTGACTGGAGTTCCTTGGCACCCGAGAATTCCA | RNA structure probing with DMS, LL_2 DMS- |
| ILMN_PCR2i18 | CAAGCAGAAGACGGCATACGAGAT **GCGGAC**GTGACTGGAGTTCCTTGGCACCCGAGAATTCCA | RNA structure probing with DMS, LL_3 DMS- |
| ILMN_PCR2i19 | CAAGCAGAAGACGGCATACGAGAT **TTTCAC**GTGACTGGAGTTCCTTGGCACCCGAGAATTCCA | RNA structure probing with DMS, HL_1 DMS+ |
| ILMN_PCR2i20 | CAAGCAGAAGACGGCATACGAGAT **GGCCAC**GTGACTGGAGTTCCTTGGCACCCGAGAATTCCA | RNA structure probing with DMS, HL_2 DMS+ |
| ILMN_PCR2i21 | CAAGCAGAAGACGGCATACGAGAT **CGAAAC**GTGACTGGAGTTCCTTGGCACCCGAGAATTCCA | RNA structure probing with DMS, HL_3 DMS+ |
| ILMN_PCR2i22 | CAAGCAGAAGACGGCATACGAGAT **CGTACG**GTGACTGGAGTTCCTTGGCACCCGAGAATTCCA | RNA structure probing with DMS, HL_1 DMS- |
| ILMN_PCR2i23 | CAAGCAGAAGACGGCATACGAGAT **CCACTC**GTGACTGGAGTTCCTTGGCACCCGAGAATTCCA | RNA structure probing with DMS, HL_2 DMS- |
| ILMN_PCR2i24 | CAAGCAGAAGACGGCATACGAGAT **GCTACC**GTGACTGGAGTTCCTTGGCACCCGAGAATTCCA | RNA structure probing with DMS, HL_3 DMS- |
| RT_15xN | AGACGTGTGCTCTTCCGATCTNNNNNNNNNNNNNNN | RNA structure probing with NAI-N3 |
| Ligation_adapter | PHO-NNNNNNNAGATCGGAAGAGCGTCGTGTAGGGAAAGAGTGT -3NHC3 | RNA structure probing with NAI-N3 |
| PCR_forward primer | AATGATACGGCGACCACCGAGATCTACACTCTTTCCCTACACGACGCT | RNA structure probing with NAI-N3 |
| PCR_reverse_index.2 | CAAGCAGAAGACGGCATACGAGAT **ACATCG**GTGACTGGAGTTCAGACGTGTGCTCTTCCGATCT | RNA structure probing with NAI-N3, sample LL1 |
| PCR_reverse_index.4 | CAAGCAGAAGACGGCATACGAGAT **TGGTCA**GTGACTGGAGTTCAGACGTGTGCTCTTCCGATCT | RNA structure probing with NAI-N3, sample LL4 |
| PCR_reverse_index.5 | CAAGCAGAAGACGGCATACGAGAT **CACTGT**GTGACTGGAGTTCAGACGTGTGCTCTTCCGATCT | RNA structure probing with NAI-N3, sample LL3 |
| PCR_reverse_index.8 | CAAGCAGAAGACGGCATACGAGAT **TCAAGT**GTGACTGGAGTTCAGACGTGTGCTCTTCCGATCT | RNA structure probing with NAI-N3, sample NAI-N3 |
| PCR_reverse_index.9 | CAAGCAGAAGACGGCATACGAGAT **CTGATC**GTGACTGGAGTTCAGACGTGTGCTCTTCCGATCT | RNA structure probing with NAI-N3, sample DMSO |
| PCR_reverse_index10 | CAAGCAGAAGACGGCATACGAGAT **AAGCTA**GTGACTGGAGTTCAGACGTGTGCTCTTCCGATCT | RNA structure probing with NAI-N3, sample in vitro |
| PCR_reverse_index12 | CAAGCAGAAGACGGCATACGAGAT **TACAAG**GTGACTGGAGTTCAGACGTGTGCTCTTCCGATCT | RNA structure probing with NAI-N3, sample LL5 |
| PCR_reverse_index13 | CAAGCAGAAGACGGCATACGAGAT **TTGACT**GTGACTGGAGTTCAGACGTGTGCTCTTCCGATCT | RNA structure probing with NAI-N3, sample LL3 |
| PCR_reverse_index14 | CAAGCAGAAGACGGCATACGAGAT **GGAACT**GTGACTGGAGTTCAGACGTGTGCTCTTCCGATCT | RNA structure probing with NAI-N3, sample HL1 |
| PCR_reverse_index15 | CAAGCAGAAGACGGCATACGAGAT **TGACAT**GTGACTGGAGTTCAGACGTGTGCTCTTCCGATCT | RNA structure probing with NAI-N3, sample HL2 |
